# Microscaled Proteogenomic Methods for Precision Oncology

**DOI:** 10.1101/796318

**Authors:** Shankha Satpathy, Eric J. Jaehnig, Karsten Krug, Beom-Jun Kim, Alexander B. Saltzman, Doug Chan, Kimberly R. Holloway, Meenakshi Anurag, Chen Huang, Purba Singh, Ari Gao, Noel Namai, Yongchao Dou, Bo Wen, Suhas Vasaikar, David Mutch, Mark Watson, Cynthia Ma, Foluso O. Ademuyiwa, Mothaffar Rimawi, Jeremy Hoog, Samuel Jacobs, Anna Malovannaya, Terry Hyslop, Karl C. Clauser, D. R. Mani, Charles Perou, George Miles, Bing Zhang, Michael A. Gillette, Steven A. Carr, Matthew J. Ellis

**Affiliations:** Broad Institute of Harvard and Massachusetts Institute of Technology, Cambridge, Massachusetts 02142, USA; Lester and Sue Smith Breast Center and Dan L Duncan Comprehensive Cancer Center, Baylor College of Medicine, Houston, TX 77030, USA; Verna and Marrs McLean Department of Molecular and Human Genetics, Baylor College of Medicine, Houston, TX, 77030, USA; Siteman Comprehensive Cancer Center and Washington University School of Medicine, St. Louis, Missouri 63110, USA; National Surgical Adjuvant Breast and Bowel Project (NSABP) Foundation, Pittsburgh, PA 15212, USA; Department of Biostatistics and Bioinformatics, Duke University Medical Center, Durham, NC, USA, 27710, USA; Lineberger Comprehensive Cancer Center, University of North Carolina at Chapel Hill, Chapel Hill, NC, USA, 27514, USA; Division of Pulmonary and Critical Care Medicine, Massachusetts General Hospital, Boston, MA 02115, USA

## Abstract

Cancer proteogenomics integrates genomics, transcriptomics and mass spectrometry (MS)-based proteomics to gain insights into cancer biology and treatment efficacy. A proteogenomics approach was therefore developed for frozen core biopsies using tissue-sparing specimen processing with a “microscaled” proteomics workflow. For technical proof-of-principle, biopsies from ERBB2 positive breast cancers before and 48-72 hours after the first dose of neoadjuvant trastuzumab-based chemotherapy were analyzed. ERBB2 protein and phosphosite levels, as well as mTOR target phosphosites, were significantly more suppressed upon treatment in cases associated with pathological complete response, suggesting MS-based pharmacodynamics is achievable. Furthermore, integrated analyses indicated potential causes of treatment resistance including the absence of ERBB2 amplification (false-ERBB2 positive) and insufficient ERBB2 activity for therapeutic sensitivity despite ERBB2 amplification (pseudo-ERBB2 positive). Candidate resistance features in true-ERBB2+ cases, including androgen receptor signaling, mucin expression and an inactive immune microenvironment were observed. Thus, proteogenomic analysis of needle core biopsies is feasible and clinical utility should be investigated.

## Introduction

Cancer proteogenomics integrates data from cancer genomics and transcriptomics with cancer proteomics to provide deeper insights into cancer biology and therapeutic vulnerabilities. Both by improving the functional annotation of genomic aberrations and by providing insights into pathway activation, this multi-dimensional approach to the characterization of human tumors has shown promise for the delineation of cancer biology and treatment options^1–5^. In addition, proteogenomics applied to patient-derived xenograft (PDX) samples has exposed potential predictive markers and mechanisms of tumor response and resistance ^3, 6, 7^.

Proteogenomics has been limited by the amount of tissue required, restricting translational research opportunities and applicability to cancer diagnostics. For example, the Clinical Proteomic Tumor Analysis Consortium (CPTAC) has required a minimum of 100mg (wet weight) of tissue from a surgical resection specimen, which typically yields several hundred micrograms of protein that provides quantitative information on >10,000 proteins and >30,000 phosphosites per sample ^8^. In these initial projects, concerns were raised related to sample heterogeneity and pre-analytical variability because RNA, DNA and protein were often isolated from separate parts of the tumor and after variable sample ischemia periods of an hour or more. For clinical diagnostics, a “microscaled” approach is required, whereby a single snap-frozen tumor-rich core needle biopsy (10 to 20 mg wet weight) must provide sufficient DNA, RNA and protein for deep-scale proteogenomic profiling that includes genome sequencing, RNA sequencing, and deep-scale mass spectrometry-based quantification of proteins and post-translational modifications. This would facilitate proteogenomic profiling of clinical biopsy specimens, including paired pre-and on-treatment analyses to determine on-target pathway inhibition and the identification of resistance mechanisms. Analysis of multiple cores could help mitigate the challenges of intra-tumoral heterogeneity.

To achieve these goals, we have developed two complementary approaches. First, methods were devised to generate high-quality DNA, RNA and protein for deep-scale DNA and RNA sequencing and proteome and phosphoproteome analysis from a single 14G core needle biopsy, with a uniform distribution of sample entering each analyte preparation protocol (Biopsy Trifecta Extraction, “BioTExt”). Second, we developed a microscaled liquid chromatography-mass spectrometry (LC-MS/MS)-based proteome and phosphoproteome analysis pipeline by executing technical refinements to reduce by more than ten-fold the amount of input required per channel in a Tandem Mass Tag (TMT)-10 or TMT11-plex experiment. This microscaled proteomics (“MiProt”) protocol requires only 25 ug peptide per sample. Using patient derived xenografts (PDX), we demonstrate that MiProt retains sufficient depth of proteome and phosphoproteome coverage when compared to standard protein input as illustrated previously ^8^.

As technical proof-of-principal, we applied these methods to a small-scale clinical study designed to test the feasibility of proteogenomic profiling before and 48-72 hours after initiating trastuzumab-based treatment for ERBB2+ breast cancer. We chose ERBB2+ breast cancer for our initial study as an example of clearly-defined oncogenic kinase-driven tumor where proteogenomic analyses and pharmacodynamic studies should provide significant insights into variability in treatment outcomes ^9, 10^. Neoadjuvant treatment with trastuzumab and pertuzumab combined with chemotherapy is the standard of care, producing pathological complete responses (pCR) in the breast and nodes in up to 75% of patients (Loibl and Gianni 2017). Further improvements in pCR and long-term remission rates depend on the identification of treatment resistance mechanisms and the ability to direct patient to effective alternatives when antibody-based targeting of ERBB2 fails.

## Results

The laboratory protocol described herein combines two microscaling methods that provide preparative and analytical approaches with the following features (Figure 1): (1) the extraction strategy (BioTExt) maximizes the yield of analytes from small clinical samples by isolating proteins for MS analysis before DNA extraction from the residual pellet; (2) the interposition of multiple 5um sections for histological analysis to provide tumor content information throughout the core biopsy; (3) a sectioning approach that ensures proteomic and genomic analyses are conducted on near identical biological replicates; (4) successful scaling of MS-based proteomics technology (MiProt) for analyses of limited amounts of biopsy derived protein.

**Figure 1:**
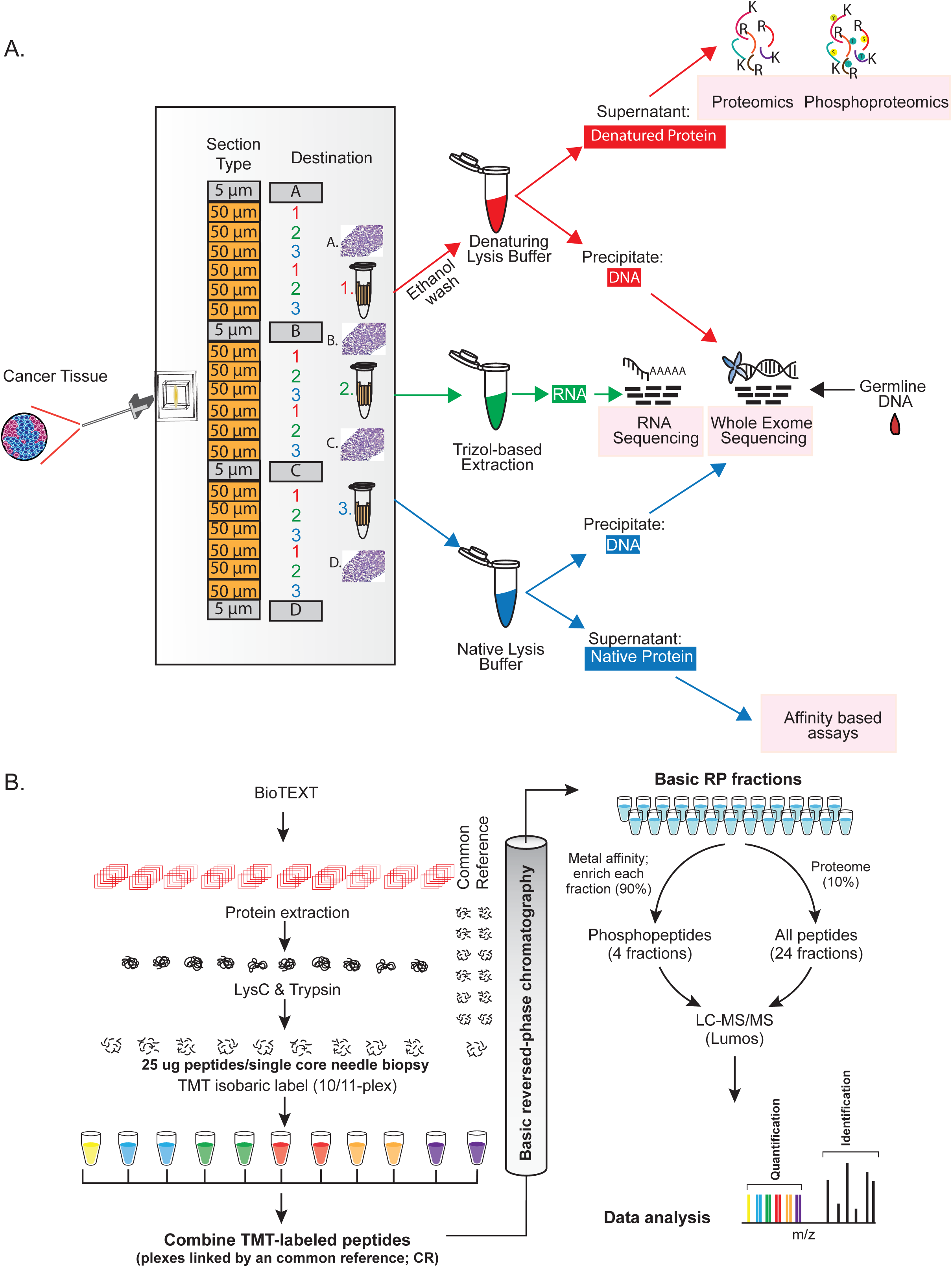
Biopsy-trifecta extraction based proteogenomics workflow. **A**. In the Biopsy Trifecta EXTraction (BioTEXT) protocol, patient derived OCT-embedded needle core biopsies are sectioned, followed up by extraction of DNA, RNA and proteins for deep-scale proteogenomics characterization and by immunohistochemistry-based imaging. **B**. The Microscaled <u>P</u>roteomics (MiProt) workflow allows deep-scale proteomics and phosphoproteomics characterization with 25ug of peptides per core-needle biopsy. MiProt uses a common reference that could be used for comparison across all samples within a single-TMT10/11 plex and across several TMT10/11 plexes spanning several core biopsies.

### Development and evaluation of the Biopsy Trifecta Extraction protocol (BioTExt)

To perform proteogenomics analyses from flash-frozen diagnostic core needle biopsies, we devised and optimized the BioTExt protocol. A single optimal cutting temperature (OCT)-embedded core biopsy was serially sectioned with alternating 50um sections transferred into 3 different 1.5ml tubes (Figure 1A). A total of six sections were transferred into each tube. To assess sample quality, 5um sections were taken before the first and after every sixth 50um section for H&E staining, with adequate quality control requiring 50% average tumor content throughout the sample. The first tube was used to extract denatured protein and DNA, the second tube was used for RNA isolation, and the third tube was used to extract native protein and DNA. The denatured protein was subsequently used for proteomic and phosphoproteomic analyses described herein, and DNA and RNA were used for genomic analysis. The native protein analyses will be described elsewhere.

### Development and evaluation of microscaled proteomics

To assess the quantity of recoverable analytes using the procedure outlined above, we applied BioTExt to several OCT-embedded core-needle biopsies collected from a total of 4 previously established breast cancer patient-derived xenograft (PDX) models: WHIM2, WHIM14, WHIM18 and WHIM20 ^11^. The yield for the sum of all six sections from a single biopsy in these PDX tumors ranged from 2.5-14 ug DNA, 0.9-2.3 ug RNA and 280-430 ug of protein. Extraction yields for the nucleic acid extractions are provided in Supplementary Figure 1A. The yields of the three analytes required a method capable of providing a deep-scale proteome and phosphoproteome despite lower analyte input. Because a wide range of needle sizes (14-22 gauge) are used to obtain diagnostic biopsies and different tumor types yield widely varying amounts of protein, a minimum of 25ug of input peptide/sample was set as the target. This amount should reasonably and consistently be obtained from six 50um curls from a needle core biopsy even in low-yield tumors or when using small biopsy gauges. BioTExt also allows additional sections from each core to be reserved for verification studies, such as targeted MS analyses once candidate proteins and phosphosites of interest have been identified or for replication of full-depth discovery analyses if required due to technical failures.

To obtain deep proteome and phosphoproteome coverage from 25ug of input peptide/sample, a tandem mass-tagging (TMT) peptide labeling approach was employed ^12^ (Figure 1B). Since the mass tags are isobaric, signals from the same peptides in each sample stack at the MS1 level, improving overall sensitivity for identification and quantification, a key advantage for the analysis of small amounts of protein. Multiplexing also increases sample analysis throughput by 10-fold relative to label-free approaches. Successful microscaling required several modifications to the bulk-optimized CPTAC workflow ^8^ to allow labeling, fractionation and analysis of low amounts of proteins. This overall method is referred to as “Microscaled Proteomics” (MiProt).

To determine if the proteomic coverage for core-needle biopsies are comparable to those obtained using a workflow optimized for bulk tumors (the Clinical Proteomics Tumor Analysis Consortium (CPTAC) workflow) ^8^, a head-to-head comparison experiment utilizing previously published breast cancer PDX models was executed ^11^, including two luminal (WHIM18 and WHIM20) and two basal-like models (WHIM2 and WHIM14) (Figure 2A). To simulate a diagnostic analysis, two needle-biopsy cores were collected from each PDX model. The cored xenograft tumors were then surgically removed for analysis of the residual bulk material. The cores were OCT-embedded, flash frozen and subjected to BioTExt followed by MiProt. The remaining bulk tumors were flash frozen and cryopulverized, followed by analysis using the original CPTAC workflow ^8, 13^. Totals of 300ug of peptides per sample were analyzed with the original and 25ug of peptides per sample with the MiProt workflow using a randomized experimental layout (Supplementary Table 1).

**Figure 2:**
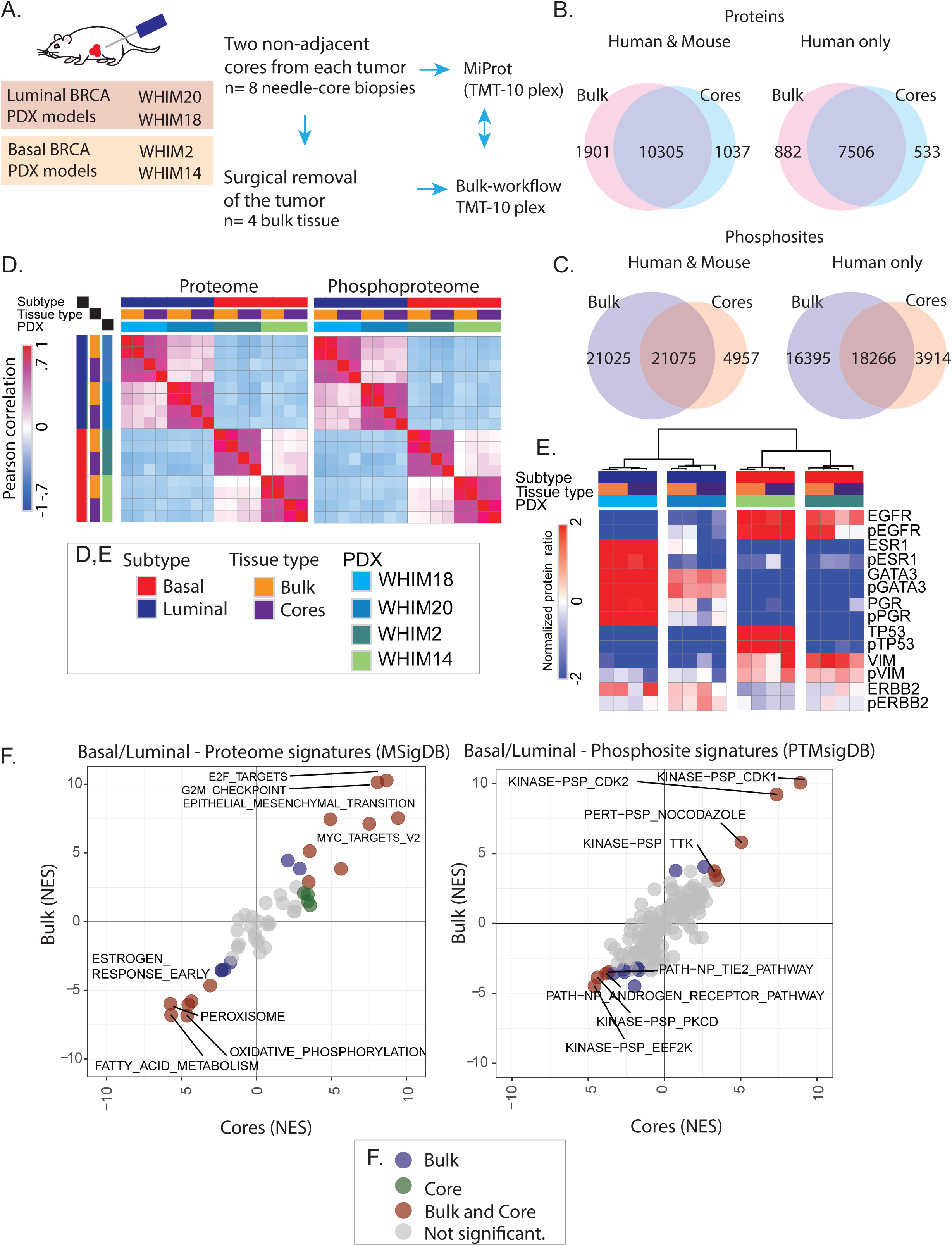
Evaluation of BioText and MiProt workflow on preclinical PDX models. **A**. Non-adjacent, core needle biopsies were collected from 2 basal and 2 luminal PDX models followed by surgical removal of tumors. Proteomic and phosphoproteomic characterization of cores were performed using the MiProt workflow, and the bulk tissue was characterized using CPTAC workflow described in Mertins, et.al. ^8^ **B** Venn-diagram showing the number of overlap between human and mouse or human proteins quantified in cores and bulk tissue. **C**. Venn-diagram shows the overlap between human and mouse or human phosphosites. **D**. Pearson correlation of TMT ratios for proteins (left) and phosphosites (right) between each sample from both cores and bulk across all 4 PDX models. **E**. The heatmap shows the TMT ratios for key differentially regulated Luminal versus Basal breast cancer associated proteins and phosphoproteins (average expression of identified phosphosites) identified across both bulk and cores experiments. **F**. Gene-centric and phosphosite-centric pathway or kinase activity enrichment analysis was performed using GSEA (MSigDB “Cancer Hallmarks”, left) and PTM-SEA (PTMSigDB, right), respectively, for Luminal-Basal differences captured in bulk (y-axis) and core (x-axis) tissue. limma derived signed Log10 p-values were used to pre-rank differential features for both GSEA and PTM-SEA analysis. The pathway/phospho-signatures that are significant in both cores and bulk are indicated in brown.

Protein and phosphosite expression were reported as the log ratio of each sample’s TMT intensity to the intensity of an internal common reference included in each plex. Both workflows identified more than 10,000 proteins, of which >7,500 were identified as human. Extensive overlap was observed between the populations of proteins identified by the two workflows (Figure 2B, Supplementary Table 2A). MiProt identified >25,000 phosphosites from each core, and these sites showed substantial overlap with those identified by high input, bulk workflows (Figure 2C, Supplementary Table 2B, Supplementary Figure 2A). The identification of over 25,000 phosphosites in the MiProt method is of particular note as this is less than a two-fold reduction in quantified sites relative to the CPTAC bulk workflow ^8^ despite using 12-fold less tumor material than the bulk workflow. While prior studies have reported relatively high numbers of proteins from small amounts of tissue material (∼4500 from small amounts of tissue material)^14^, the very large number of phosphosites we obtained using just 25 ug of peptide/sample has not been described previously. There was a high correlation of TMT ratios between replicates of bulk tumors and between replicates of cores across all 4 PDX models for both the proteomics and phosphoproteomics data (Figure 2D, Supplementary Figure 2B). In addition to a high degree of overlap in protein and phosphosite identities, expression was also highly correlated (R> 0.65) between cores and bulk for individual PDX models, as can be visualized by the close juxtaposition of core and bulk samples from the same PDX model upon unsupervised hierarchical clustering (Supplementary Figure 2C).

Expression profiles of key basal and luminal markers, showed some degree of heterogeneity, although, showed a comparable trend overall, between bulk and core analysis (Figure 2E) at the levels of both the proteome and phosphoproteome for all PDX models except for WHIM20, where phosphorylated EGFR, phosphorylated PGR and ESR1 protein showed reduced expression in cores relative to bulk, suggesting that there may be some heterogeneity in this particular PDX model as shown previously ^11^. By contrast, ERBB2, a breast cancer marker that should not be highly expressed in these clinically ERBB2-(no ERBB2 amplification) cases showed more uniform expression across the different PDX models. Overall, cores provided proteomics data that yielded results consistent with those obtained from global expression profiles from bulk tissue.

To address whether differentially regulated pathways and phosphosite-driven signaling in luminal versus basal subtypes were captured by the microscaled workflow, pathway-level and kinase-centric analyses were applied to the bulk and core sample data. Single-sample gene-set enrichment analysis (ssGSEA) was applied to proteomics data, and post-translational modifications set enrichment analysis (PTM-SEA) to the phosphoproteomic data ^15, 16^. The luminal-basal differences captured by bulk tissue analysis were highly correlated with differences detected using cores for both protein and phosphosite expression (Figure 2F, Supplementary Table 2C, D). Of note, the data recapitulates previously observed luminal-basal differences, which provided a quality metric for the proteomics dataset both for cores and bulk tissue ^2, 6^. The same conclusion was reached in bulk versus core comparisons performed on the normalized TMT protein ratios for individual PDX models (Supplementary Figure 2D). Despite identifying ∼40% fewer phosphorylation sites, most of the differential Luminal-Basal kinase signatures identified in the bulk tissue were captured by MiProt (Figure 2F, right).

### Application of microscaled proteogenomic analyses to clinical core biopsies from patients treated for ERBB2+ locally advanced breast cancer

The effectiveness of the BioTExt and MiProt analyses in PDX models encouraged the application of these methods to clinical tumor samples acquired in the context of a small-scale ERBB2+ breast cancer study (Discovery protocol 1 (DP1); NCT01850628). This study was designed primarily to investigate the feasibility of proteogenomic profiling before and immediately after initiating trastuzumab-based treatment for ERBB2+ breast cancer. Patients with a palpable breast mass diagnosed as ERBB2 positive breast cancer by a local laboratory were treated at the physicians’ discretion, typically with trastuzumab in combination with pertuzumab and chemotherapy. The regimens included docetaxel or paclitaxel, the former often combined with carboplatin. The protocol (https://clinicaltrials.gov/ct2/show/study/NCT01850628) was designed to study acute treatment perturbations by accruing OCT-embedded core needle biopsies before and 48 to 72 hours after treatment (referred to pre-treatment and on-treatment, respectively, throughout the text).

As shown in the REMARK (Reporting Recommendations for Tumor Marker Studies) ^17^ diagram (Supplementary Figure 3), core biopsy samples were available from 19 patients. Proteogenomic analysis could be conducted on samples from 14 patients as five cases had low tumor content biopsies. For the included samples, the analyte yield varied across different cores, but the lower-range yields of DNA, RNA and protein (0.4ug, 0.2ug, and 45ug respectively) were sufficient to demonstrate the suitability of the optimized extraction protocol for clinical biopsy specimens (Supplementary Figure 1B). Protein, and RNA when available, was also analyzed for on-treatment cores from 10 patients, with analysis of duplicate pre- and on-treatment cores achieved in four of the patients, and of triplicate cores in one patient (Figure 3A). In total, 35 cores were analyzed. Tumor and germline whole-exome sequencing was performed using DNA from a single baseline core for all 14 patients. DNA isolated from cores using BioTExt yielded target coverage comparable to that from genomic DNA isolated from blood (generated using standard organic extraction techniques), indicating there was no compromise in DNA quality when a protein-sparing technique was used (Supplementary Figure 4A). Transcriptome analysis was successful for 30 cores with available RNA, corresponding to 11 of the 14 patients, and TMT 11-plex-based proteomics analysis was performed for all 35 cores.

**Figure 3:**
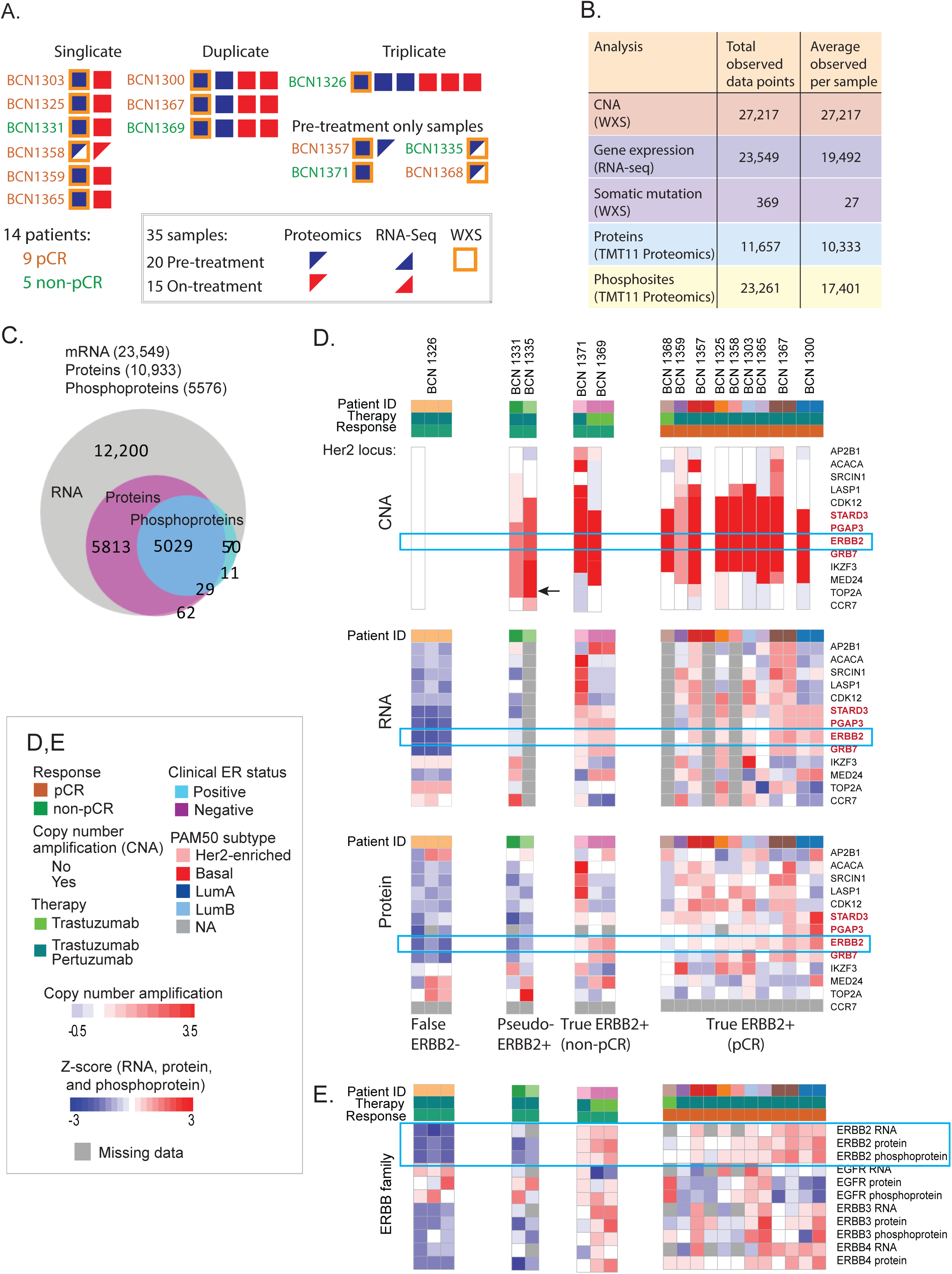
Application of microscaled proteogenomics to the “Discovery protocol 1” clinical trial (DP1), a small scale trial focused on trastuzumab-based neoadjuvant chemotherapy. **A**. Overview of proteogenomics samples obtained from pre- and on-treatment core biopsies from the DP1 clinical trial. Each block indicates the data obtained from a separate core. **B**. Microscaled proteogenomics achieves a high level of proteogenomics depth for the DP1 core needle biopsies. Table summarizing total proteogenomics coverage and numbers of mutated genes for all samples and average coverage across all analyzed cores is shown on the left. **C**. The Venn-diagram shows the overlap between all genes identified across RNA-seq, proteomics and phosphoproteomics. **D**. Heatmap summarizing proteogenomics features of ERBB2 amplicon and adjacent genes at the level of CNA, RNA and Protein expression. The set of genes in red make up the core of the ERBB2 amplicon and showed consistently high copy number amplification, RNA, and protein levels in all of the pCR cases (True ERBB2+ pCR set on the right) and in BCN1371 and 1369 (True ERBB2+ non-pCR set) but significantly lower protein levels in BCN1326 (False ERBB2+) and BCN1331 and BCN1335 (Psuedo ERBB2+ set). The arrows point to the amplified TOP2A gene and protein in BCN1335. **D**. The heatmap at the bottom shows corresponding Z-scores of RNA, protein, and phosphoprotein expression of ERBB1-4 across all 14 patients. ERBB3 protein and phosphoprotein and ERBB4 protein levels were also significantly lower in BCN1326, BCN1331, and BCN1335 than in the set of pCR cases.

On average, we obtained copy number information on >27,000 genes, measured mRNA transcripts for >19,000 genes, and quantified >10,000 proteins and >17,000 phosphosites from each individual patient sample, with a large overlap of gene identification across different datasets (Figure 3B, Supplementary Table 3, 4). This was equivalent to the depth obtained in previous large-scale breast cancer proteogenomics efforts based on bulk tissues, with the exception of the phosphoproteome coverage, which achieved about half of the number of sites previously reported for tumor bulk-level characterization ^2, 18^. For 13 out of the 14 cases, ERBB2 amplification was confirmed by exome sequencing along with amplifications and mutations in a range of genes previously implicated by the TCGA and ICGC breast cancer studies (Supplementary Figure 4B) ^19, 20^, including mutations in TP53 in 6 out of the 15 patients, PIK3CA missense mutations in BCN1371 and BCN1325, and a somatic nonsense mutation in BRCA1 in BCN1335. We also observed a similar overall pattern of chromosomal amplifications and deletions between patients in this clinical trial and the subset of ERBB2+ tumors from TCGA, including a high frequency of previously described amplifications of chromosomes 1, 8, and 20 (Supplementary Figure 4B) ^20^. The median gene-wise Spearman correlation between mRNA and protein across patient cores was 0.38, consistent with previous bulk-focused proteogenomics studies (Supplementary Figure 4C) ^1, 2, 4, 18^. In addition, co-expression networks derived from MiProt protein expression better predicted KEGG pathway function than those derived from mRNA expression for a similar proportion of pathways as previously reported for the published CPTAC breast cancer cohort ^21^ (Supplementary Figure 5A). Deep-scale proteogenomic analysis of duplicate and pre- and on-treatment cores from the same patient allowed an assessment of BioTExt sample processing reproducibility.

Unsupervised hierarchical clustering based on 500 most-variable genes resulted in all duplicate cores clustering together at the level of mRNA, protein and phosphosite expression with the exception of samples from case BCN1365, where pre- and on-treatment profiles did not cluster together at the level of phosphosite expression (Supplementary Figure 5B). In summary, the data generated using microscaled methods was of comparable quality to previous bulk tissue-focused proteogenomics reports and yielded observations that were consistent with the expected biology for tumors diagnosed as ERBB2+.

### Proteogenomic analysis of the ERBB2 locus suggests false positive clinical diagnoses

All the patients in this study were locally diagnosed as ERBB2+ based on standard fluorescence *in situ* hybridization (FISH) and immunohistochemistry (IHC)-based approaches. A pathological Complete Response (pCR) occurred in 9/14 cases (64%), but 5 patients had residual cancer at surgery (non-pCR). To probe the possibility that some of the non-pCR cases were due to misassignment of ERBB2 status, proteogenomic analysis of the region of chromosome 17q spanning the ERBB2 locus and adjacent genes was performed (Figure 3D). Most obviously, exome sequencing of BCN1326 did not show amplification of ERBB2 or other nearby genes (Figure 3D, upper panel) and exhibited markedly lower levels of ERBB2 RNA (Figure 3D, middle panel) and protein expression (Figure 3D, lower panel) than pCR cases, suggesting a false positive (False ERBB2+). Additionally, expression levels from genes immediately flanking ERBB2 (STARD3, PGAP3 and GRB7, highlighted in red in Figure 3D) were lower than pCR cases. BCN1331 and BCN1335 may represent a more subtle form of false positivity. While these samples showed a gain of ERBB2 copy number, ERBB2 protein levels remained low, similar to BCN1326. BCN1335 also showed greater absolute amplification of TOP2A than of ERBB2 (see black arrow Figure 3D upper panel), and the TOP2A protein was markedly over-expressed compared to all other cases (Figure 3D lower panel, of note the RNA analysis failed in this sample). This suggests that TOP2A was the more likely driver in this case. Levels of STARD3, PGAP3 and GRB7 both for RNA (BCN1331) and protein (BCN1331 and BCN1335) were also low, indicating that the amplicon may not drive sufficient ERBB2 expression for treatment sensitivity, which, we refer to as “Pseudo ERBB2 positive”. When comparing these three false positive samples as a group with the nine “true” ERBB2 positive pCR cases, both the arithmetic mean for STARD3, ERBB2 and GRB7 protein log TMT ratios and the protein log ratios of each gene separately were significantly lower in the proposed false ERBB2 positive cases (mean: p=0.0114, STARD3: p=0.0255, ERBB2: p=0.0073, GRB7: p=0.0399). Protein levels of ERBB2 dimerization partners ERBB3 and ERBB4, as well as phospho-ERBB3, were also significant under-expressed in all three proposed false positive-samples when compared to the pCR cases (ERBB3 protein: p=0.0097, ERBB3 phosphoprotein: p=0.0318, ERBB4 protein: p=0.0131) (Figure 3E). In contrast, protein and phosphoprotein levels of EGFR, the remaining dimerization partner of ERBB2, does not appear to be correlated with ERBB2 levels and was high in the non-pCR samples, suggesting that EGFR homodimers may play a driver role in signaling when ERBB2 is low ^22, 23^. Thus, BCN1335 and BCN1331 represent “pseudo” ERBB2 positive cases, i.e. false ERBB2 positive cases for which proteogenomic evidence reveals insufficient ERBB2 expression to attain pCR despite gains in ERBB2 gene copy number. A central immunohistochemistry analysis indicated that ERBB2 was 1+ in BCN1326 and 2+ in BCN1335 and BCN1331, all the pCR cases were assigned 3+ staining (Supplementary Figure 6A). In contrast to the three ErbB2 low-expression cases, BCN1371 and BCN1369 were both non-pCR cases despite exome-confirmed ERBB2 amplification, ERBB2 RNA and protein expression similar to pCR cases with IHC 3+ ERBB2 staining and over-expression of STARD3, PGAP3, GRB7, ERBB3 and ERBB4 (Figure 3D). These two cases therefore represent examples of true positive cases with intrinsic therapeutic resistance. Supporting the quantitative potential of microscaled proteomics, the samples with an IHC score of 3+ in a central assay showed significantly higher levels of ERBB2 expression than tumors scored 1+ or 2+ (p=0.00013). Parallel reaction monitoring (PRM) was also deployed as an orthogonal label-free protein quantification method on the same samples with an excellent correlation (R=0.92, p value=0) between the TMT and PRM-based MS approaches (Supplementary Figure 6B-D).

### Proteomics and phosphoproteomic analyses of acute on-treatment samples demonstrates pharmacodynamic monitoring for response prediction is achievable

The clinical study was primarily designed to test the feasibility of proteogenomics analysis to identify early markers for responsiveness to ERBB2-directed monoclonal antibody therapy Proteomics, Phosphoproteomics and RNAseq (for comparison) was therefore conducted on pre- and on-treatment core biopsies for nine patients with pCR and three patients without pCR. Differential treatment-induced changes were not observed at the RNA level and trended but did not reach significance at the ERBB2 protein level (Figure 4A; while the ERBB2 protein levels showed significant reduction in pCR cases (p=0.031), the p-value for the comparison of this reduction between pCR and non-pCR cases was 0.067). However, greater downregulation of ERBB2 phosphoprotein (mean of all ERBB2 phosphosites) levels after 48-72 hours in pCR cases than in non-pCR cases was observed (two sample sum rank test, p=0.017; Figure 4A). To explore these data further, *limma* ^24^, a more advanced statistical method specifically designed for differential expression analysis of small sample size studies, was employed. (Supplementary Table 5). Differential ERBB2 RNA expression was again not seen for any comparison (Supplementary Figure 7). However, there was significant pCR-specific downregulation for both ERBB2 protein (limma: p=0.002 for pCR, p=0.63 for non-pCR, and p=0.029 for pCR vs. non-pCR) and phosphoprotein levels (p=0.000014 for pCR, p=0.88 for non-pCR, and p=0.0086 for pCR vs. non-pCR) (Supplementary Table 5). More importantly, differential analysis of individual phosphorylation sites revealed pCR-specific significant downregulation of several phosphosites on proteins from the pathway, including sites on ERBB2 and SHC1, an adaptor that binds to ERBB2 ^25^ (Figure 4B). Most of the significant changes that were at least 2-fold and that affected the ERBB2 pathway were observed in the site-level phosphoproteomics data (Figure 4B; Supplementary Figure 7).

**Figure 4:**
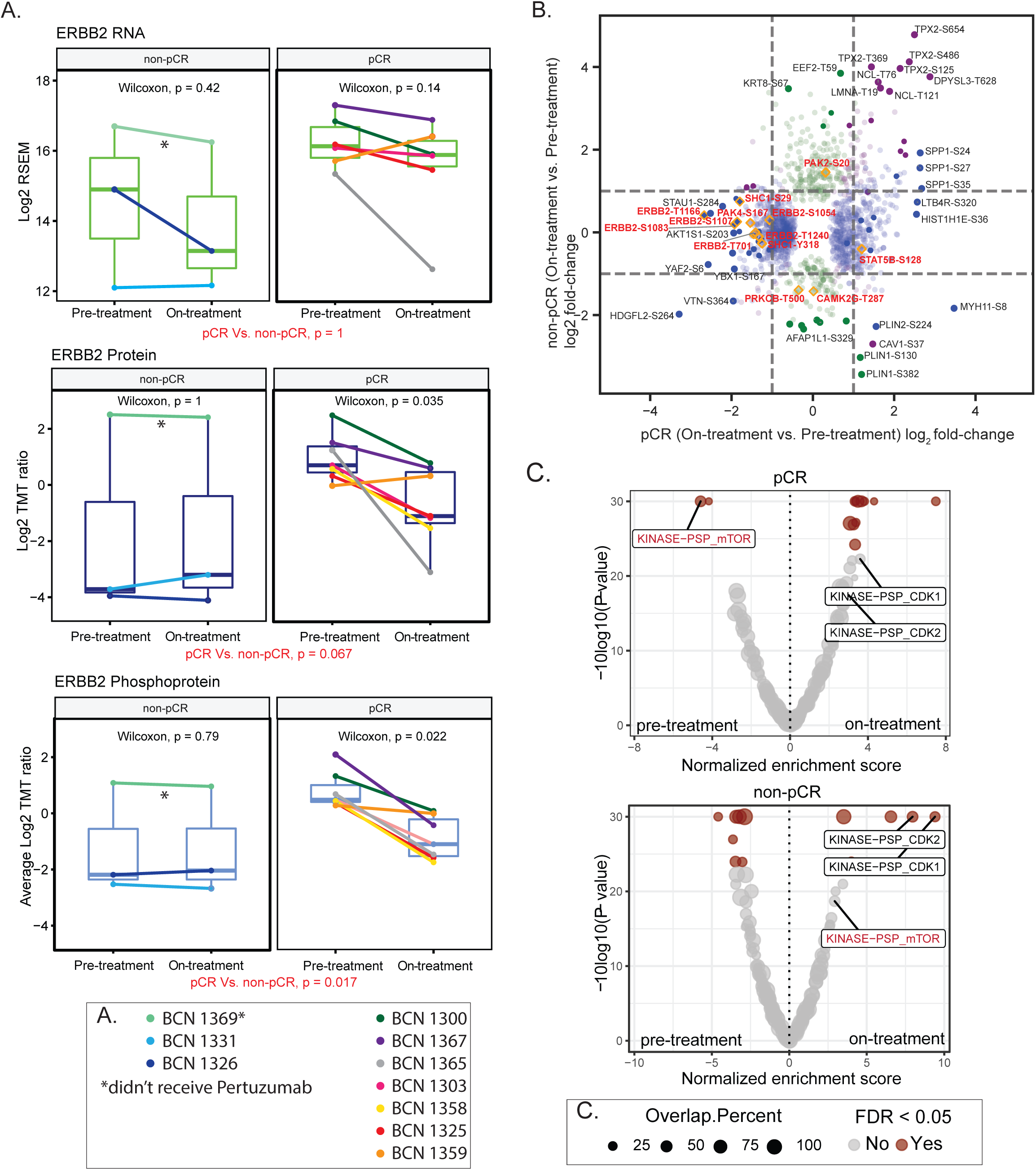
Dual anti-ERBB2 therapy results in downregulation of ERBB2 and mTOR signaling in cases with pCR. **A**. Effect of anti-ERBB2 treatment on ERBB2 RNA, protein, and phosphoprotein levels for each patient with on-treatment data. P-values for paired Wilcoxon signed rank tests for on-treatment vs. pre-treatment ERBB2 expression for each group. The pCR vs. non-pCR p-values are derived from Wilcoxon rank sum tests comparing log2 fold changes of on-treatment to pre-treatment levels from pCR patients to those from non-pCR patients. For patients with multiple cores, the mean expression value was used. **B**. Scatter plot showing differential regulation of phosphosites before and after treatment in pCR and in non-pCR cases. Shown are the on-treatment vs. pre-treatment log2 fold changes in non-pCR (y-axis) vs. the log2 changes in pCR samples (x-axis) for phosphosites with p-value <0.05 by limma analysis of differential expression in either group. Blue and green circles indicate phosphosites in pCR and non-pCR, respectively that show significant differential regulation in either group alone. Purple circles indicate significantly regulated phosphosites in both sets of patients. The orange diamond outlines highlight phosphosites on proteins in the KEGG ErbB signaling pathway (hsa04012). The transparency of each point reflects it’s significance after BH-adjustment (adjusted p<0.05 is solid, and more transparent points have higher adjusted p-values). **C**. PTM-SEA was applied to the signed Log10 p-values from *limma* differential expression analysis of on- vs. pre-treatment phosphosite levels from pCR cases (upper panel) and non-pCR (lower panel). The volcano plot shows the Normalized Enrichment Scores (NES) for kinase signatures. Red circles indicate signatures with significant FDR (<0.05).

Given the well-understood kinase signaling cascades downstream of ERBB2 ^22^, a recently published tool for pathway analysis of phosphosites, PTM-SEA, was applied to the phosphoproteomics data ^15^. This program uses a manually curated post-translational modification site database, PTMsigDB (https://github.com/broadinstitute/ssGSEA2.0), to estimate the activity for phosphoproteomics signatures resulting from chemical or genetic manipulation of a pathway or for kinases by analyzing signatures for target substrates with validated biochemical evidence. Figure 4C shows significant phosphosite signatures for the comparisons tested (on- vs pre-treatment changes in pCR only and in non-pCR only) (Supplementary Table 6). Supplementary Figure 8 shows a heatmap of phosphoproteome driven signatures that were significantly differentially regulated (FDR <0.05) upon treatment in either of the two groups. While the inferred activities of CDK1 and CDK2 kinases (KINASE−PSP_CDK1, KINASE−PSP_CDK2) were upregulated in the non-pCR patients, downregulation of mTOR activity (KINASE-PSP_mTOR) was most prominent exclusively in pCR cases upon treatment. This comparison of pre- and on-treatment samples thus suggests that *in vivo* downregulation of mTOR signaling, downstream of ERBB2, during treatment leads to a more favorable response.

### Pathway enrichment analysis reveals variability in the tumor immune microenvironment and other putative response features in individual non-pCR cases

To explore candidate biological processes that may contribute to inadequate response to therapy in non-pCR cases, RNA, protein and phosphoprotein outlier analyses on data from each pre-treatment core from the non-pCR cases with respect to the set of pre-treatment pCR cores was performed. Specifically, Z-scores were calculated for each gene/protein in a given individual non-pCR core relative to the distribution established from all of the pre-treatment pCR cores. The Z-scores of ERBB2 protein expression in non-pCR cases were consistent with the observations noted above; ERBB2 RNA, protein and phosphoprotein levels in patients BCN1326, BCN1331 and BCN1335 were outliers with negative Z-scores while ERBB2 expression in patients BCN1369 and BCN1371 lied within the normal distribution of the pCR cases (Figure 5A, Supplementary Figure 9). Z-scores derived from the outlier analysis for each of the data points (RNA, proteome and phosphoproteome; see Supplementary Table 7A) were used for single sample Gene Set Enrichment Analysis (ssGSEA). Figure 5B highlights a subset of immune-centric and oncogenic signaling pathways that showed differential enrichment in the non-pCR cases. The expanded list is available as Supplementary Table 7B. Consistent, significantly enriched pathway-level differences across replicate cores and multiple data types from a single patient was observed (Figure 5B), confirming that microscaled proteogenomics data obtained from cores in a clinical setting yield reproducible results. Interestingly, distinct biological pathways showed differential enrichment in each of the individual non-pCR cases relative to the pCR class (Figure 5B, Supplementary Table 7B).

**Figure 5:**
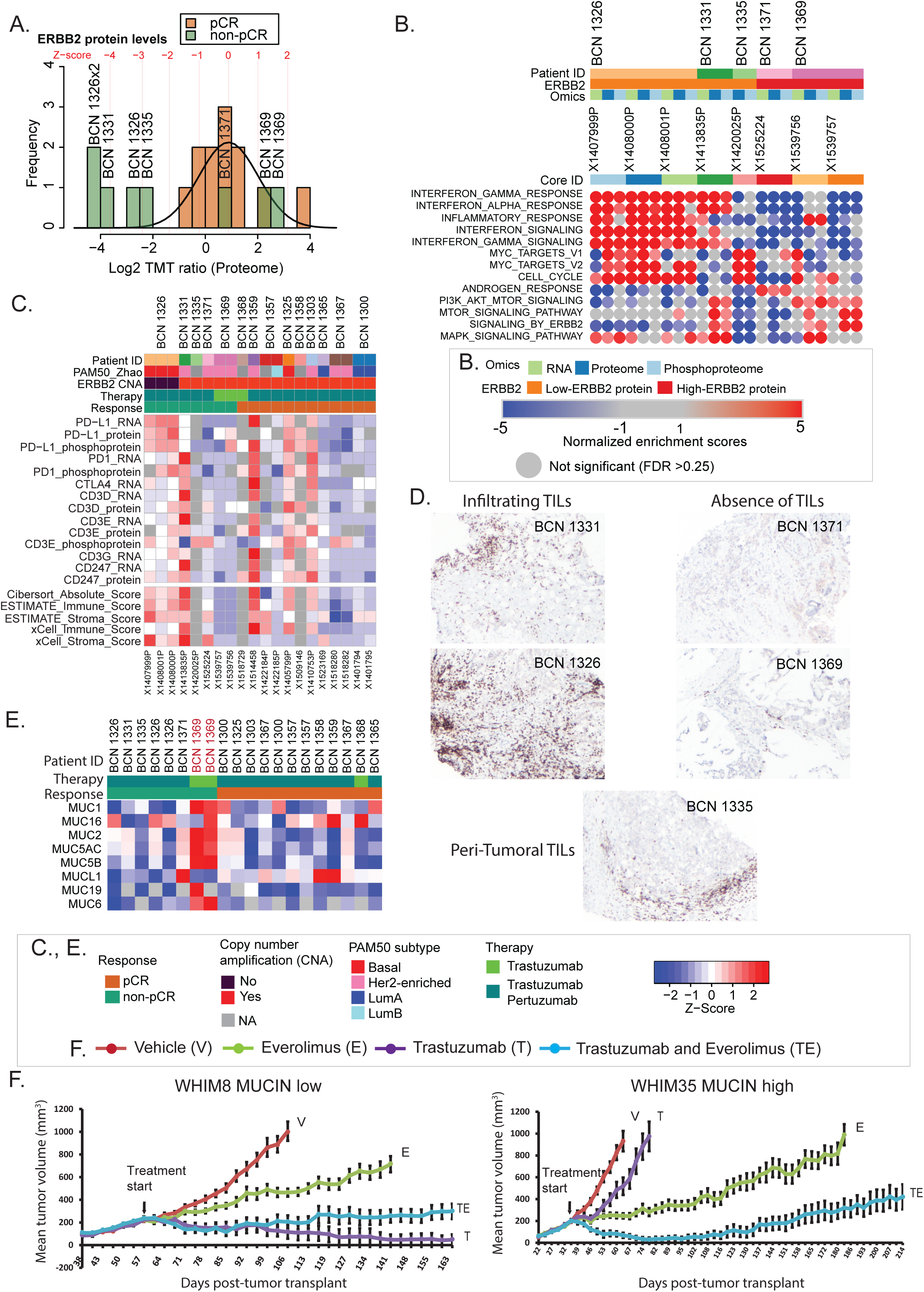
Proteogenomics analysis of baseline untreated samples suggest diverse “candidate” mechanisms of resistance in non-pCR cases. **A.** Outlier analysis was performed to identify differentially regulated mRNA, proteins or phosphoproteins in each pre-treatment sample from non-pCR cases relative to the set of pre-treatment samples from all pre-treated pCR cases. Shown is the ERBB2 protein distribution across all patients; brown and green bars indicate the frequencies for each protein level bin in non-pCR and pCR, respectively. The line shows the normal distribution of pCR samples from which the Z-score for each non-pCR sample was derived. Corresponding Z-scores levels are indicated in red. **B.** Heatmap showing normalized enrichment scores (NES) from single sample Gene Set Enrichment Analysis (ssGSEA) of outlier Z-scores from non-pCR cases. Shown are a subset of differentially regulated pathways with false-discovery rate less than 25% (FDR <0.25). **C.** Heatmap showing expression levels of key immune-checkpoint and T-cell marker (CD3) genes and of RNA based immune and stroma scores from ESTIMATE, Cibersort, and xCell. **D.** Photomicrographs showing anti-CD3 immunohistochemical staining profiles of non-pCR cases (original magnification: 20x). **E.** Heatmap showing Mucin protein expression across all pre-treated patients. **F.** WHIM8 and WHIM35 PDX models were treated with vehicle, trastuzumab, everolimus or the combination of trastuzumab and everolimus. The graph shows the mean-tumor volume at several timepoints after tumor implantation and subsequent treatment, and error bars show standard error of mean.

Of the complex patterns revealed by differential pathway analysis immune-related and interferon signaling pathways showed consistent upregulation across the data sets in samples from two of the three cases with lower expression of ERBB2, BCN1326 and BCN1331. In contrast, these pathways showed variable downregulation for the remaining non-pCR cases. To further explore these findings, the expression of T cell receptor (CD3 isoforms and CD247) and immune checkpoint (PD-L1, PD1, and CTLA4) genes were analyzed and immune profiles from the RNA-seq data using established tools were generated (Cibersort, ESTIMATE, and xCell). Examination of immune profile scores and of expression of T cell receptors and targetable immune checkpoint regulators supported the presence of an active immune response in BCN1326 relative to other samples (Figure 5C, Supplementary Figure 10). Similarly, immune profile scores also indicated that BCN1331 had an activated immune microenvironment, and PD1 RNA expression was higher in this patient than in any other case (Figure 5C). The five non-pCR cases were stained for the pan T-cell marker CD3 to validate these proteogenomic findings (Figure 5D). Consistent with the active immune microenvironment (Figure 5C), both BCN1326 and BCN1331 demonstrated tumor T-cell infiltration. In contrast, a predominant peri-tumoral or “immune-excluded” inflammatory reaction was observed in BCN1335 and a complete paucity of T-cells (immune-desert) was observed in the two resistant proteogenomically confirmed ERBB2+ cases, BCN1371 and BCN1369, consistent with the lack of immune signaling documented in Figure 5B.

Other variable differential features in resistant cases included PI3K/AKT/mTOR and MAPK signaling, all of which represent potential therapeutic opportunities. ERBB2 pathway activation in BCN1331 is unexpected given the very low level of ERBB2 protein but could be explained by expression of EGFR/pEGFR (Figure 3E). MYC targets were consistently upregulated at the protein and phosphoprotein levels in BCN1326 and BCN1335 and the androgen response pathway was upregulated at all levels in BCN1371. Consistent with the elevated AR signaling observed in BCN1371 (Figure 5B), this tumor exhibited histologic features of an apocrine cancer with intensely eosinophilic cytoplasm and AR expression (Supplementary Figure 11 middle and lower panel). Interestingly, BCN1331 also expressed AR by IHC (Figure 5D) without activation of an androgen response signature or apocrine features (Supplementary Figure 11), consistent, with the disconnect between AR expression and AR signaling in breast cancer noted by others ^26^. Also, for patient BCN1371, we did not see significant upregulation of PI3K signaling (Figure 5B) despite PIK3CA mutation (E545K), consistent, with the disconnects between PIK3CA mutation and effects when signaling was assessed by reverse phase protein array (RPPA) ^19, 27^. Table 1 summarizes the proteogenomic features observed for each non-pCR case.

**Table 1.** A table summarizing proteogenomic features and potential mechanisms of resistance for tumors from non-pCR cases

| Case Number | RNAseq-PAM50 Subtype and ER status | HER2 Amplicon Structure by WES | "Druggable Mutations" | Differential Pathways (relative to pCR cases) | TILs and Immune Microenvironment Proteomics | Treatments Received |
| --- | --- | --- | --- | --- | --- | --- |
| BCN 1326 | Basal ER+ | No copy gain (not amplified)<br>Low ERBB2 |  | MYC UP<br>Mitosis UP<br>Interferon signaling UP | Infiltrating TILs<br>PDL1 high<br>Phospho-PD1 high | Doc, CP, T, P |
| BCN 1331 | HER2-E ER+ | Broad lower-level copy gain<br>Low ERBB2 (pseudo-amplified?) |  | ERBB2/3<br>MAPK UP<br>PI3K UP<br>mTOR UP<br>Interferon signaling UP | Infiltrating TILs<br>Phospho-PD1 high | Doc, CP, T, P |
| BCN 1335 | RNA failure ER+ | Amplicon driving TOP2A not ERBB2<br>Low ERBB2 (pseudo-amplified?) | BRCA1, R1788>stop<br>VAF=30% | MYC UP<br>Mitosis UP | Peri-tumoral TILs | Doc, CP, T, P |
| BCN 1371 | HER2-E ER- | Amplified | PIK3CA E545K<br>VAF=73.5% | AR transcription UP<br>ERBB2/3 DOWN<br>MAPK DOWN<br>PI3K DOWN<br>mTOR DOWN | No TILs | Doc, CP, T, P |
| BCN 1369 | HER2-E ER- | Amplified |  | ERBB2/3 UP<br>MAPK UP<br>PI3K UP<br>mTOR UP<br>MUCIN expression UP | No TILs | Pac, T |

### ERBB2+ Patient derived xenograft (PDX) models provide a setting to explore candidate trastuzumab resistance mechanisms and therapeutic alternatives

To further explore therapeutic resistance pathways in the proteogenomic data, association analyses between patient-centric RNA, protein and phosphoprotein outliers and the published literature were performed (Supplementary Figure 12). For each gene or protein, the terms “breast cancer” and “resist OR recur” were used to query the text from freely accessible article abstracts and keywords in the PubMed database to search for previously studied associations between the outlier genes and breast cancer resistance. Genes with highest filtered PubMed citation counts include well-studied proteins such as ESR1, BRCA1/2, TP53, EGFR, and AKT1 in addition to ERBB2 (Supplementary Table 8). As expected, ERBB2 was among the most prominent negative protein and phosphoprotein outliers in BCN1326, BCN1331, and BCN1335 that were associated with the keyword “resistance” (Supplementary Figure 12). Furthermore, TOP2A also stands out as being strongly associated with “resistance” from other outliers with high protein and phosphoprotein levels in BCN1335, the psuedo-ERBB2+ patient for whom the amplified locus appears to be driving TOP2A rather than ERBB2 expression (Supplementary Figure 12; Figure 3D). However, the most prominent proteomics outlier for patient BCN1369, MUC6, was not associated with citations containing the keyword “resistance” (Supplementary Figure 12). Nonetheless, multiple mucin family members were outliers with high protein expression specifically in this patient, two of which had citations associated with “resistance” (Supplementary Figure 12). The consistently high levels of mucin protein expression in patient BCN1369 are clearly discernible in the heatmap shown in Figure 5E. This observation is notable because mucin expression has been proposed to mask ERBB2 epitopes and prevent trastuzumab binding, as shown previously in cell lines ^10, 28, 29^.

Since therapeutic hypotheses cannot be explored directly in non-pCR patients, the foregoing hypotheses about potential mechanisms of resistance are entirely speculative. To build an approach whereby potential resistance mechanisms could be explored experimentally, a published proteogenomic analysis was analyzed to determine of any of features of the resistant tumors in the DP1 study were phenocopied in ERBB2+ patient-derived xenografts (PDX) ^6^. This analysis focused on two ERBB2+ PDX models, WHIM8 and WHIM35 both of which were shown to be responsive to lapatinib, a small molecular inhibitor against ERBB2, indicating they were true ERBB2+ cases ^6^. Interestingly, WHIM35 has high expression of mucin proteins compared to WHIM8 (Supplementary Figure 13A) indicating this case was a phenocopy of BCN1369 (Supplementary Figure 13B). Consistent with cell line-based studies reported in the literature ^30–33^ that proposed that mucin expression might inhibit trastuzumab-mediated response, trastuzumab induced tumor regression in WHIM8 but not in WHIM35 (Figure 5F). Drawing from the observation that BCN1369 also exhibited elevated PI3K-Akt-mTOR signaling (Figure 5B), together with a recent report that showed mTOR mediated MUC1 induction in multiple breast cancer cell lines ^30^, these PDX models were additionally treated with the small molecule mTOR inhibitor everolimus. Everolimus in combination with ERBB2-targeted therapy induced significant regression (Figure 5F) in the otherwise trastuzumab-resistant WHIM35 model, providing support for mucin overexpression as a “candidate” resistance mechanism that could potentially be overcome by treatment with a small molecule inhibitor of ERBB2 or downstream targeting of mTOR activity.

## Discussion

In this study, we achieved deep-scale multi-omics profiling of core needle biopsy material obtained in a clinical setting using the combined, integrative, tissue-sparing “BioText” approach described in the manuscript and applied this optimized microscaling methodology to a small cohort of breast cancer patients treated with chemotherapy and anti-ERBB2 therapy. Recently, Humphrey et al also described deep-scale phosphoproteomics analyses of cultured cell line lysates by label-free, single shot LC-MS/MS^34^. This study obtained ∼14,000 phosphosites when starting with 200ug of protein and ca. 4000 phosphosites using 25 ug of input protein. Prior efforts at proteomic and phosphoproteomic analysis of core needle biopsies of tumor tissue using “one-shot” data-independent analysis ^14, 35, 36^ or off-line SCX fractionation combined with a super-SILAC approach quantified ∼2,000–5,000 proteins and ∼3800 phosphorylation sites per core. Importantly, none of these prior studies carried out genomic analyses on the same set of samples. In contrast, our workflow provided deep-scale genomic, proteomic and phosphoproteomic analysis, identifying more than 11,000 proteins and 25,000 phosphorylation sites in PDX tissue and >17,000 in clinical tumor needle cores for integrative multi-omics analyses. The alternating tissue sectioning approach provided exceptional control over sample quality, reduced sampling bias and ensured sample consistency across the multi-omics analysis. Some of the improvement in depth that we demonstrate is undoubtedly due to the use of newer generation LC-MS/MS instrumentation. Nonetheless, an optimized multiplexing protocol enabled us to achieve this depth from as little as 25 ugs of peptides per core, rendering the pipeline viable for material obtained attained from a typical 14 to 22-gauge clinical biopsy needle.

We illustrate the potential utility of these microscaled methods by applying them to a breast cancer clinical study, a key component of which was the collection of on-treatment samples 24-48 hours after anti-ERBB2 therapy was initiated. This allowed an assessment of the immediate effects of inhibiting the ERBB2 pathway and potentially provides an early time point to determine whether a patient is likely to experience a pCR. Despite the small cohort size, we detected statistically significant downregulation of ERBB2 protein and phosphosite levels and of a phosphosite signature for downstream mTOR targets in pCR patients. Of the 7 (out of 21 total ERBB2 sites identified in Supplementary Table 4) phosphosites from ERBB2 with complete data across the cohort, all showed downregulation to varying extents in the pCR cases (Supplementary Table 5). Of the 21 sites identified, only two have been characterized in detail in cell lines (see www.phosphosite.org). These are pY-1248, a known auto-activation site ^37^, and pT-701, which may serve as a negative feedback site ^38^, although their in-vivo roles are largely unexplored. The role of downregulation of ERBB2 phosphorylation in response to treatment is complicated by the observed downregulation of ERBB2 protein levels, but from a biomarker perspective these are secondary questions that do not negate the primary conclusion that we were able to make a valid pharmacokinetic observation. We would point out that hitherto essentially all our understanding of the complex signaling properties of ERBB2 arise from experimental systems not, as we illustrate here, from patients under anti-ERBB2 treatment. Most importantly, our ability to resolve complexity in this setting to assess inhibition of ERBB2 signaling is also revealed by downregulation of a signature of target sites for mTOR, a kinase activated downstream of ERBB2, specifically in pCR patients (Figure 4C). Since treatment with other inhibitors may not directly affect the protein or phosphorylation levels of their targets, the observation of an effect on downstream signaling in the phosphosite data provides critical support for the efficacy of our microscaling approach to assess response in future studies.

An initial proteogenomic focus on ERBB2 is readily justified given the biological variability within tumors designated ERBB2 “positive”. The testing guidelines are designed to offer as many patients anti-ERBB2 treatment as possible, even though it is recognized that this “catch all” approach likely includes a number of true-negative cases that do not benefit from these treatments ^39^. Our analysis is not intended to be definitive or clinically actionable as the sample size is small and our pipeline is research-based. However, our preliminary analyses suggest three classes of resistance mechanisms to ERBB2-directed therapeutics can be defined. False-positives are exemplified by case BCN1326. In retrospect, this case was initially diagnosed by FISH but ERBB2 protein was not over-expressed when re-analyzed using standard IHC (IHC 1+). Three independent pretreatment and three post treatment biopsies were analyzed, which helps rule out tumor heterogeneity as a likely cause of the misdiagnosis. The second class of potential misclassification is “pseudo-ERBB2 positive”. As represented by cases BCN1331 and BCN1335, there was evidence for amplification of ERBB2, but proteogenomic evidence suggests that ERBB2 is not a strong driver including: a) lower levels of ERBB2 protein and phosphoprotein compared to pCR cases; b) low expression from other genes within the minimal ERBB2 amplicon (STARD3, PDAP3 and GRB7); and c) a paucity of expression of dimerization partners ERBB3 and ERBB4. Our successful validation of ERBB2 levels using single shot parallel reaction monitoring hints at a more efficient approach than the TMT multiplex assay that ultimately could form the basis of a clinical assay (supplementary Figure 6). The third resistance class demonstrated lack of pCR despite proteogenomic evidence for true ERBB2 positivity. Here proteogenomic analysis provided candidate mechanisms of resistance to consider, such as the upregulation of mucin proteins, active androgen signaling or the lack of an antitumor immune response.

We emphasize our purpose herein is not to make definitive clinical conclusions, but to illustrate the wide range of resistance biologies that microscaled proteogenomics methodologies can reveal, thus promoting further investigation. We certainly acknowledge that the therapeutic alternatives suggested in this pilot study require considerable further study. For example, for BCN1335 the proteogenomic profile (both DNA and protein) suggests that TOP2A is a more likely driver, with higher amplification and protein expression than ERBB2. Here, ERBB2 was on the “shoulder” of the amplicon giving rise to the potential misdiagnosis. The treatment of ERBB2+ breast cancer has moved away from anthracyclines ^40^, but, perhaps in cases such as this, doxorubicin could be reconsidered ^41^. The comprehensive nature of the proteogenomic data allows efficient exploration of multiple causes for treatment failure at the level of pathway activity, illustrated by androgen receptor signaling in BCN1371 (Figure 5B) and mucin expression in BCN1369 (Figures 5E). For both examples, there is prior evidence for a role in resistance to trastuzumab but persistent controversy regarding the clinical actionability of these proposed mechanisms ^29, 42, 43^.

The PDX experiments we describe are designed to illustrate how proteogenomic analyses can be used to identify individual PDX that “phenocopy” hypothetical resistance mechanisms observed clinical specimens thus promoting preclinical investigation^3, 6^. Here we were able to identify a mucin-high (WHIM35) and a mucin-low (WHIM8) ERBB2+ pair of PDX tumors suitable for exploring alternative treatments for mucin positive, trastuzumab-resistant cases. Pathway analyses of the mucin-positive clinical case BCN1369 indicated strong mTOR activity at the RNA, protein and phosphoprotein levels, suggesting that a therapeutic intervention with everolimus, an FDA-approved rapamycin-based mTOR inhibitor for ER+ advanced breast cancer ^44^ could provide an effective treatment in the setting of mucin-driven resistance. Subsequent therapeutic modeling confirmed everolimus increased trastuzumab efficacy in WHIM35 but not in the mucin-negative WHIM8 PDX, where trastuzumab alone was effective and everolimus ineffective. We would point out that from a proteogenomic standpoint mucin expression alone is likely not adequate to identify the correct population for a prospective clinical trial, as the establishment of true ERBB2+ status and a high mTOR signature could also be important.

Another important feature of the microscaled proteogenomic analysis presented herein is the ability to assess the immune microenvironment. This has become a critical aspect of breast cancer diagnostics with the approval of the PDL1 inhibitor atezolizumab in PDL1+ advanced TNBC^45^. PDL1 IHC is used as a predictive biomarker for atezolizumab, but the optimal approach to the analysis of the immune microenvironment remains under investigation^46^. BCN1326 and BCN1331, examples where the diagnosis of ERBB2 positive status was challenged by proteogenomic analysis, displayed proteomic evidence for PDL1, phospho-PD-L1, and phospho-PD1 expression, consistent with the infiltrating TIL patterns that were observed. Thus, in the future, PD1/PDL1 assessment by proteogenomics could be considered for prediction of PDL1/PD1 antibody efficacy.

While the microscaled proteogenomic methods were deployed here in the context of a clinical trial in breast cancer, they are patently extensible to any other solid tumor. The analyses described are not designed for clinical use, although potentially the time required to go from needle core biopsy to actionable results (2 to 4 weeks) is similar to next generation DNA and RNA sequencing. Analysis time could be reduced with automation of sample processing, the use of faster instrumentation and orthogonal gas phase fraction such as FAIMS ^47–49^. Furthermore, the protocol as presented can be readily adapted for use as a diagnostic tool by redirecting some of the denatured protein obtained using the BioTExt procedure to PRM assays developed for targets delineated in larger clinical discovery datasets, and, as illustrated for ERBB2 (Supplementary Figure 6).

In conclusion, our study provides the methodology for proteogenomic analysis of core biopsy material from cancer patients. The small cohort size prevents any definitive conclusions regarding the clinical utility however we have demonstrated that the identification of relevant proteogenomic features in core biopsies is a feasible exercise. We can now seek definitive clinical conclusions through analyses involving larger numbers of patients.

## Supplementary Figure legends

**Supplementary Figure 1.**
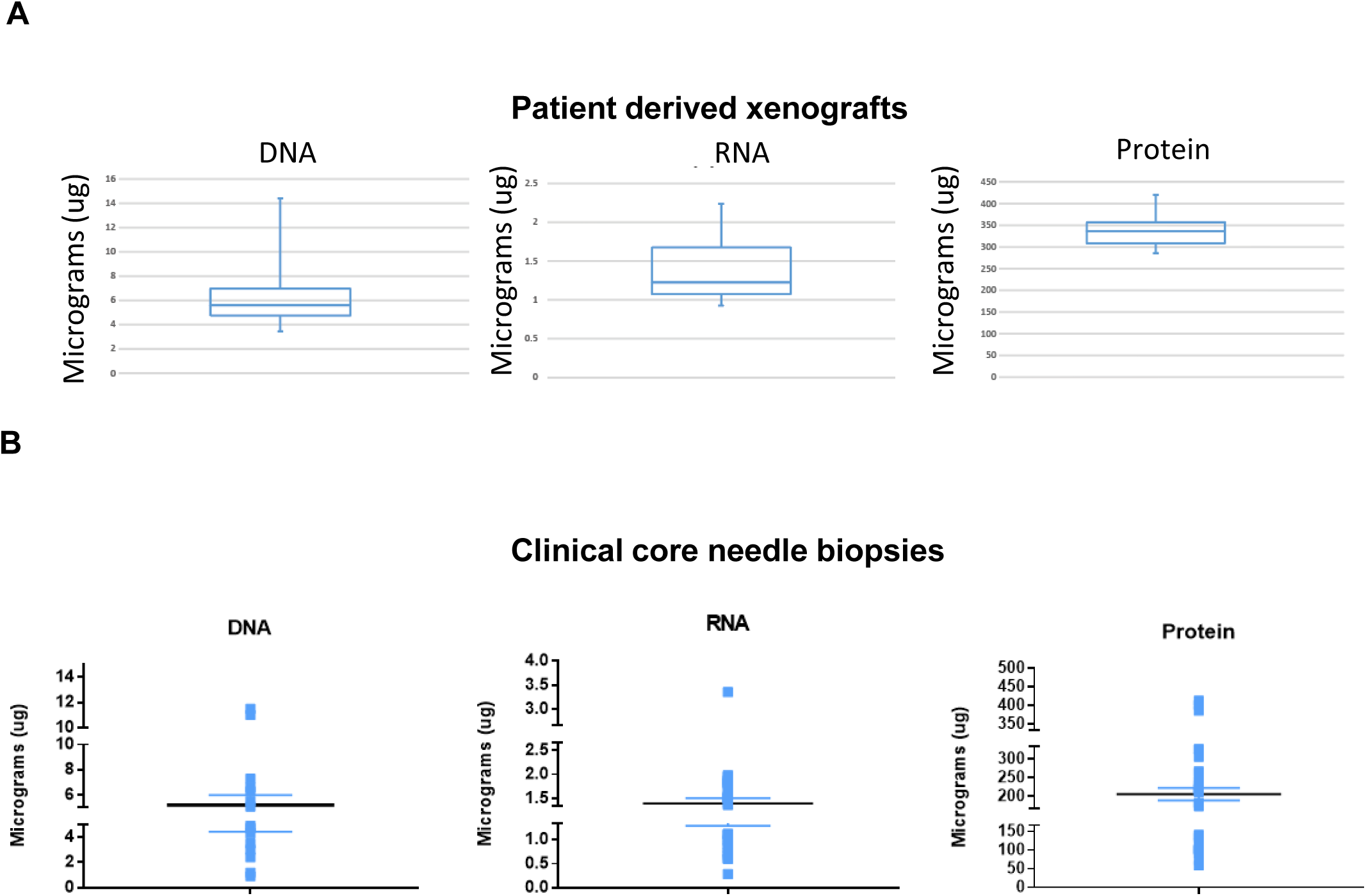
DNA, RNA and protein yields from core needle biopsy processed using BioTExt. **A**. Box plot showing DNA, RNA and Protein yields from a total of 8 core needle biopsies from 4 PDX Models: WHIM4, 14, 18 and 20. B. Box and scatter plots showing DNA, RNA and protein yields from all core needle biopsies that were processed. Samples with no yield were excluded. Error bars represent standard error of mean (SEM)

**Supplementary Figure 2.**
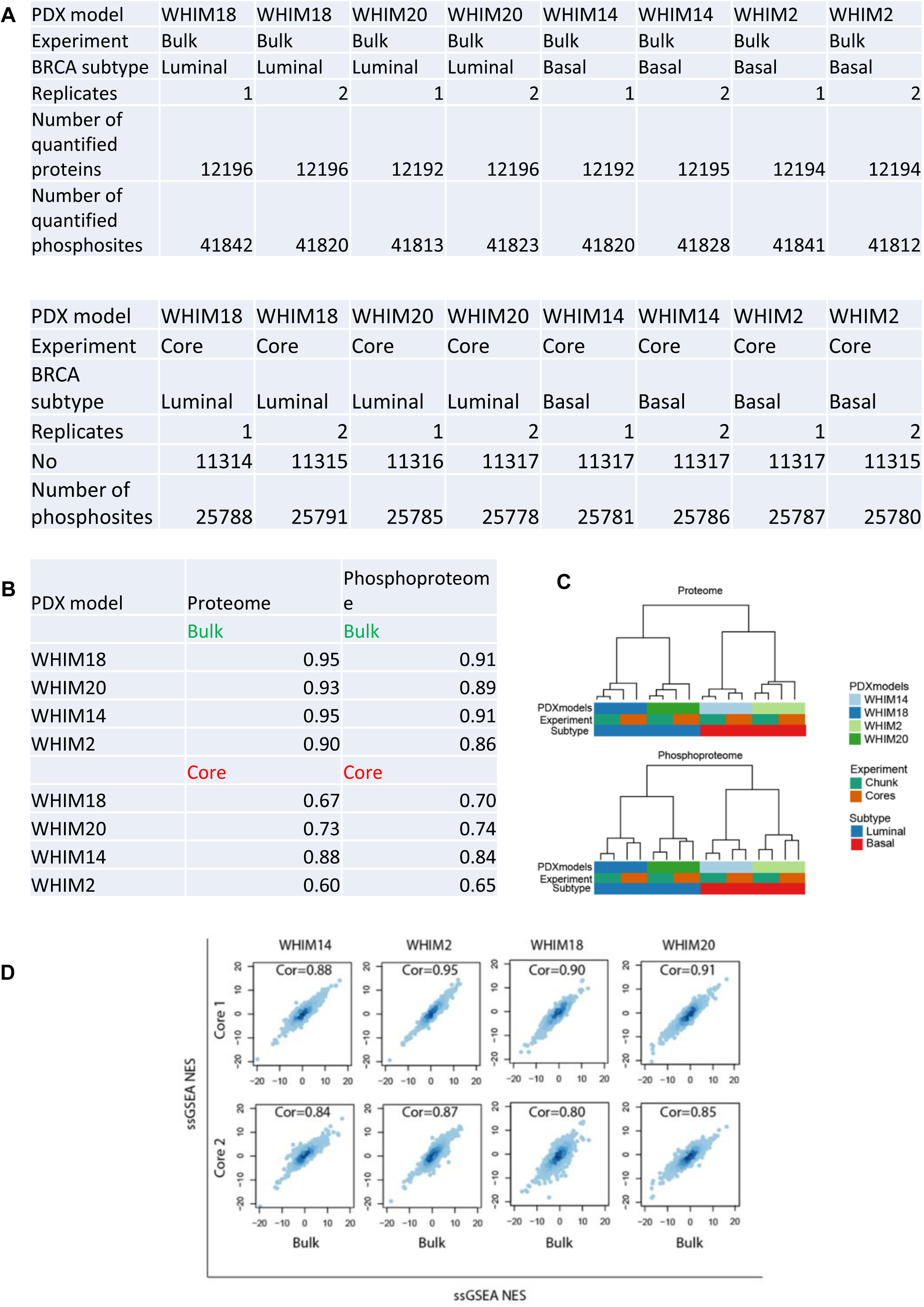
Comparison of proteomics and phosphoproteomics dataset from tumor bulk and core samples. **A.** The table shows the number of proteins and phosphosites quantified in the bulk tissue (upper panel) and non-adjacent (lower panel) cores from 4 WHIM PDX models **B.** The table lists the Pearson correlation between replicate bulk and non-adjacent cores for each of the PDX models **C**. Unsupervised hierarchical clustering (1-Pearson) of normalized TMT protein and phosphosite ratios. **D**. ssGSEA was performed on normalized TMT protein ratios obtained from cores and bulk. Scatter plot shows ssGSEA normalized enrichment scores (NES) between cores and bulk tissue for individual PDX models.

**Supplementary Figure 3.**
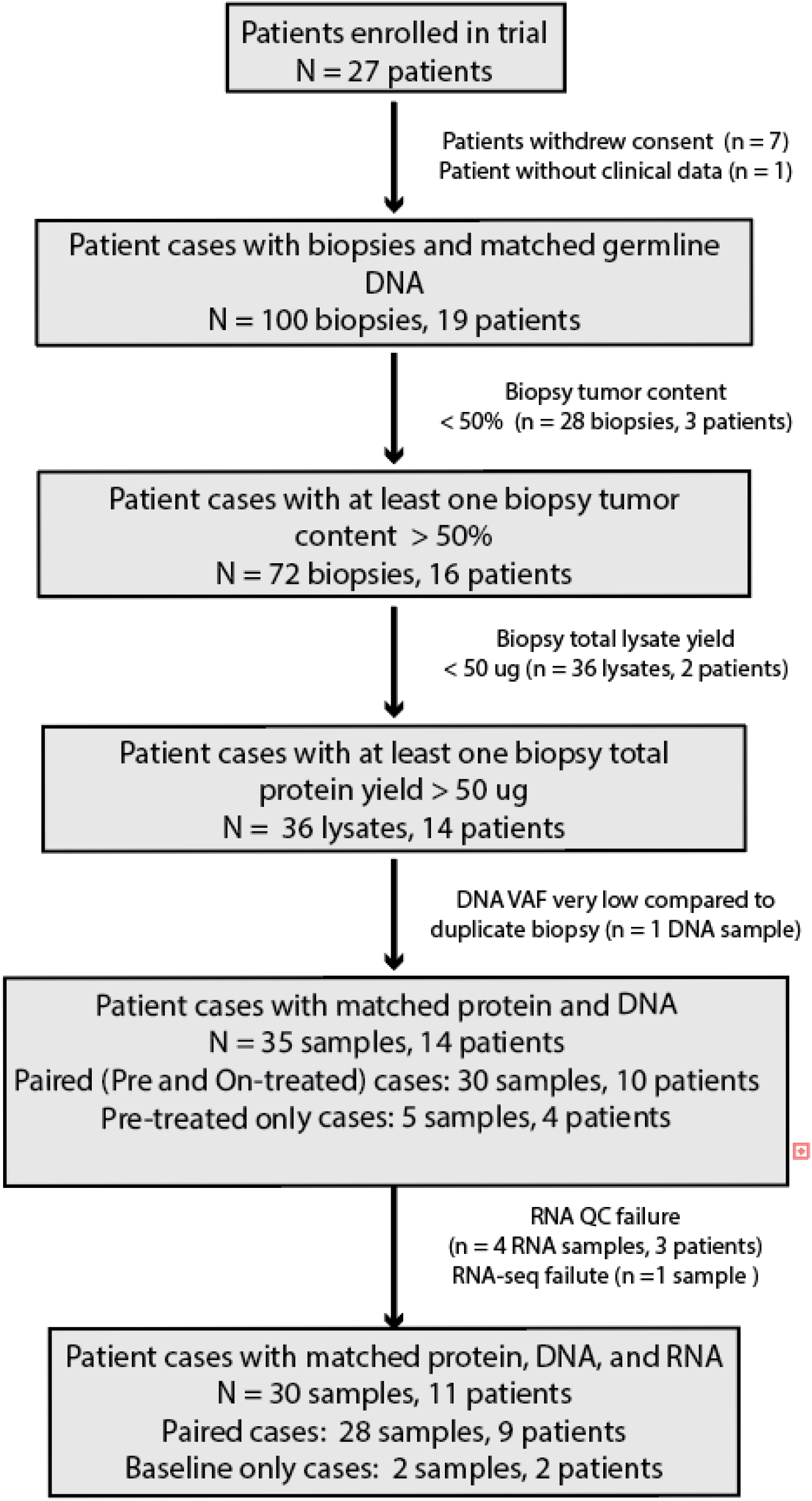
Reporting Recommendations for Tumor Marker Prognostic Studies (REMARK) diagram. The flowchart shows the number of patients enrolled in the trial and reasons for their exclusion from the proteogenomic analysis when applicable.

**Supplementary Figure 4.**
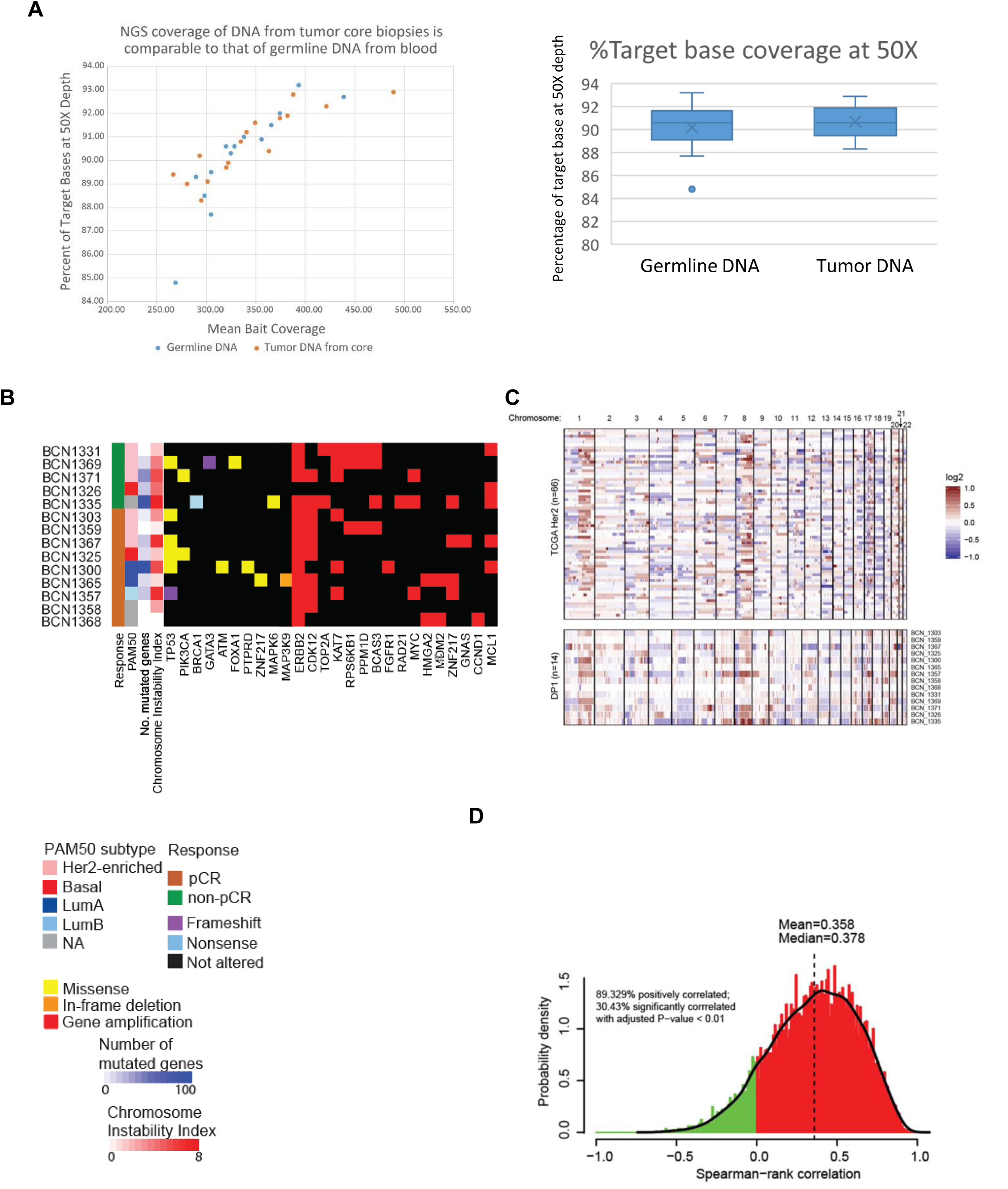
Proteogenomics features of the clinical cores. **A.** The right panel shows scatter plot between percentage of target base at 50X sequencing depth and mean bait coverage for whole-exome sequencing and the left panel shows comparable box plot distribution of percentage of target based at 50X sequencing depth for DNA isolated from blood versus tumor using BioTExt. **B.** Heatmap summarizing genomic alterations of breast cancer associated genes in tumors from 14 patients. **C**. The copy number landscape of ERBB2+ samples from TCGA (top) resembles the landscape from this study (bottom). Plots show log2 ratios of chromosome segment copy number in tumor DNA relative to normal DNA for each patient (rows) from each cohort. **D**. Distribution of gene-wise Spearman correlations between RNA and Protein as observed using the BioTExt pipeline. Red and green indicate all positively and negatively correlated genes respectively.

**Supplementary Figure 5.**
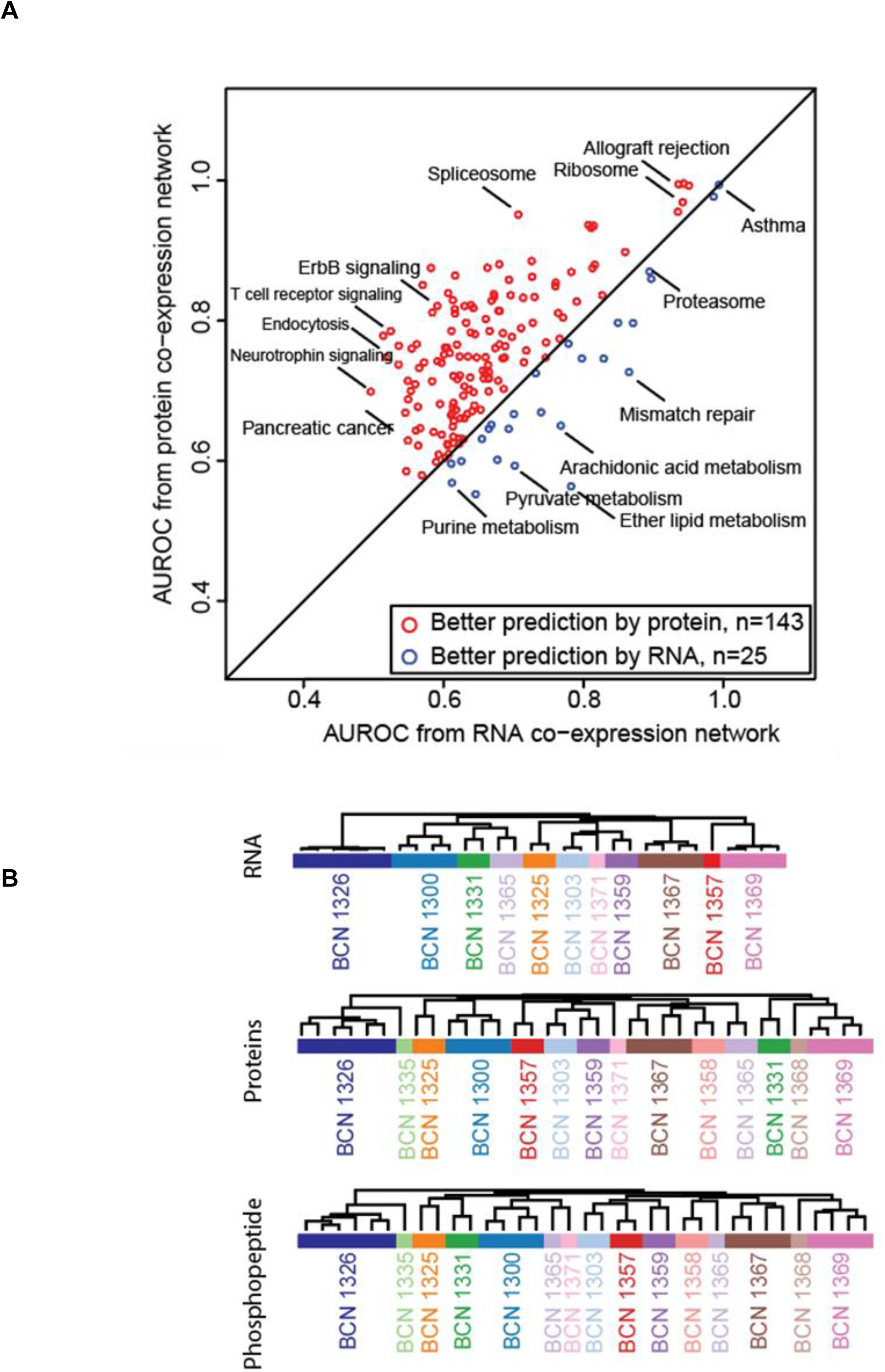
Functional prediction from co-expression networks derived from and unsupervised hierarchical clustering of samples from clinical core proteogenomics data. **A**. Co-expression networks derived from microscaled proteomics data predict function more consistently than co-expression networks derived from the RNA data. Red and blue circles indicate functional categories (KEGG pathways) predicted by co-expression networks derived from protein and mRNA expression data, respectively. **B**. Core needle biopsies from the same patients cluster together based on the top 500 most variable features in each dataset.

**Supplementary Figure 6.**
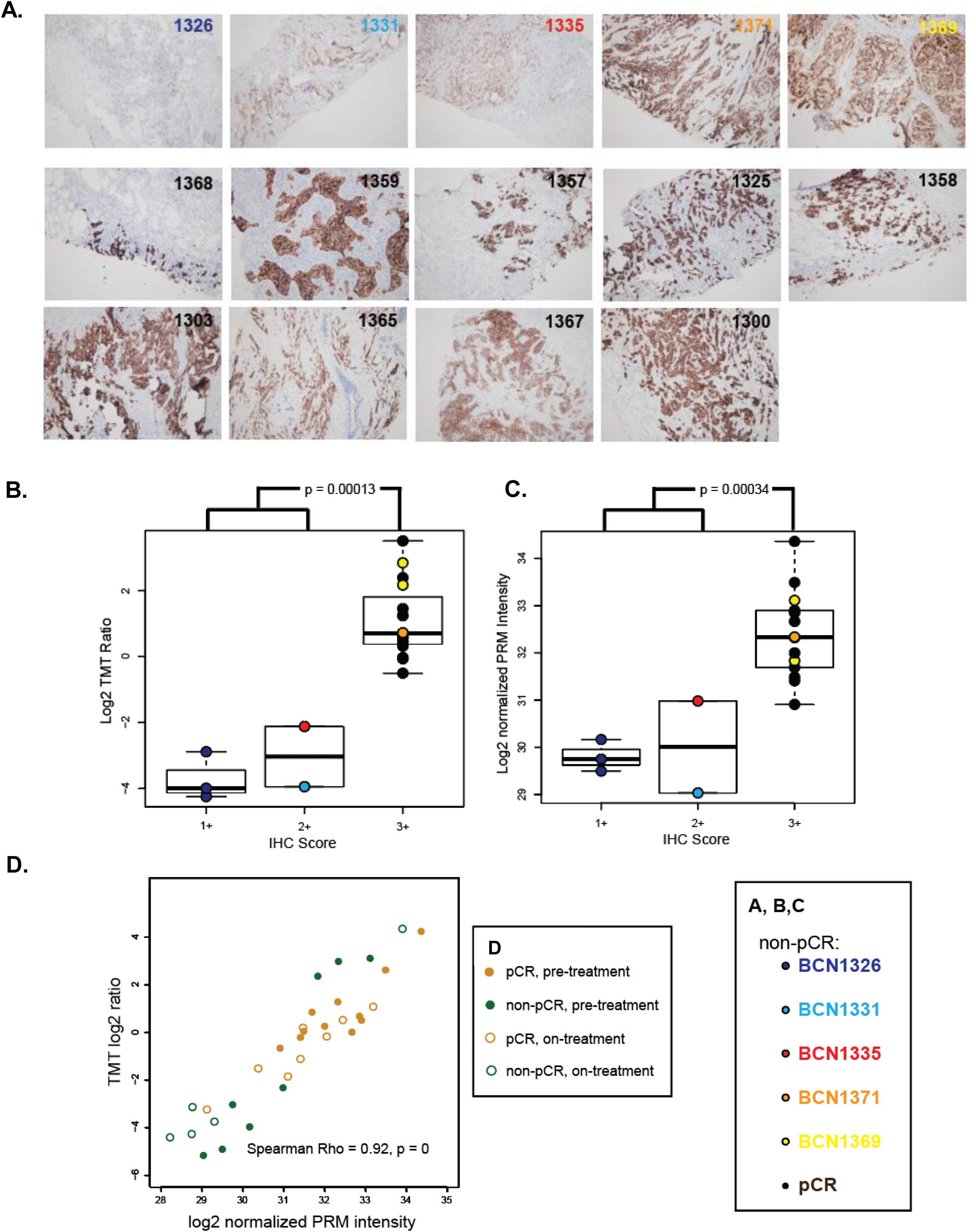
Validation of ERBB2 levels. A. ERBB2 (HER2) immunohistochemistry (IHC) on sections from all 14 patients. A. Photomicrographs showing ERBB2 IHC staining profiles of all pCR cases at 200X. B. Box plot showing ERBB2 IHC scores and ERBB2 protein levels C. Box plot showing ERBB2 IHC scores and ERBB2 protein levels as measured by parallel reaction monitoring (PRM). D. Scatter plot showing correlation between ERBB2 protein abundance measured using TMT and PRM based protein quantification.

**Supplementary Figure 7.**
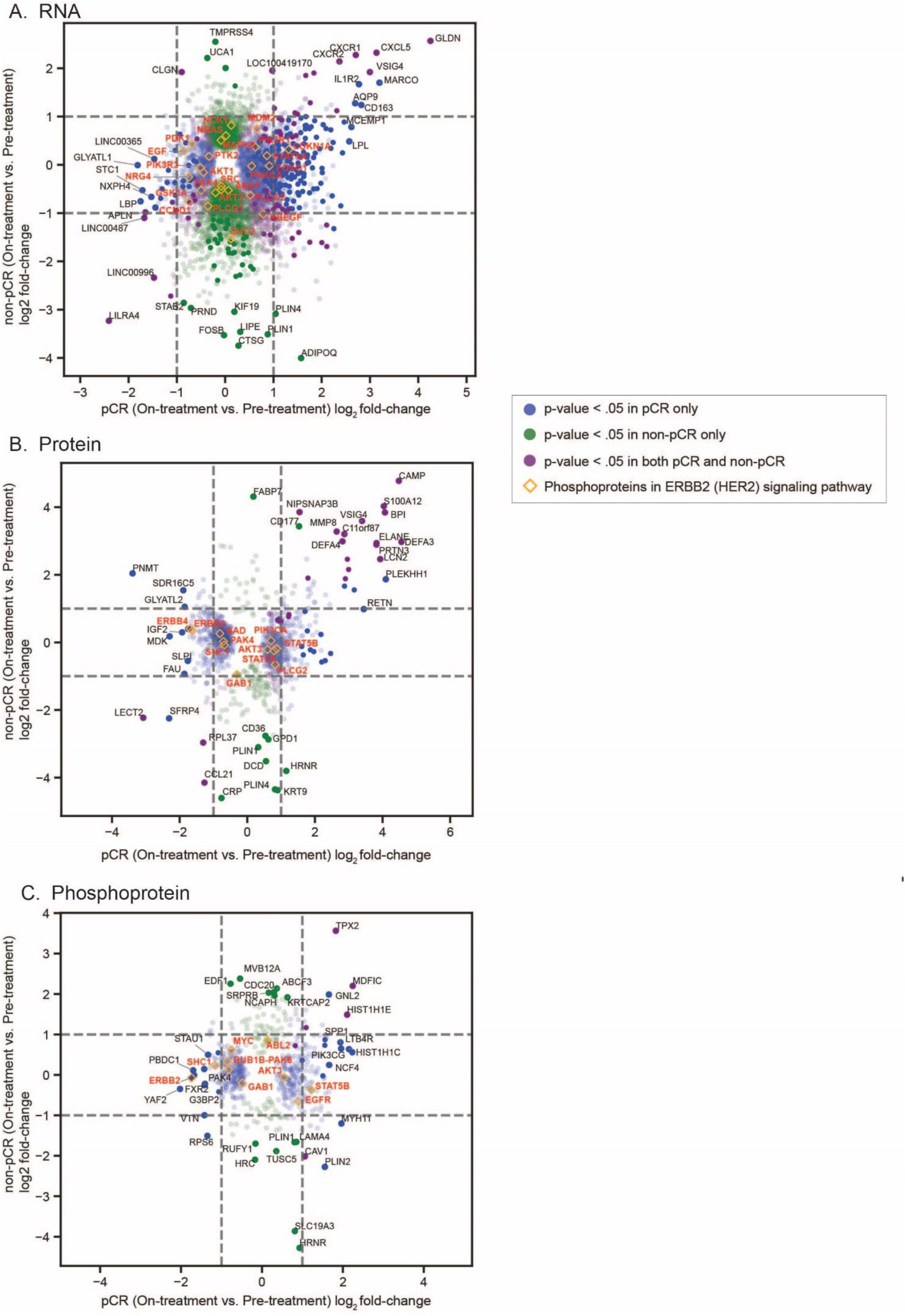
Scatter plots of response to treatment (on-treatment vs pre-treatment) in non-pCR (y-axis) vs. pCR patients for RNA, proteins, and phosphoproteins (mean of phosphosites). Shown are log2 ratios from *limma* linear modeling of differential expression for genes with p<0.05 in each set of patients. Genes from the ERBB signaling KEGG pathway (hsa04012) are highlighted in orange. The level of transparency of each point reflects it’s significance after BH-adjustment (adjusted p<0.05 points are completely opaque, and more transparent points have higher adjusted p-values).

**Supplementary Figure 8.**
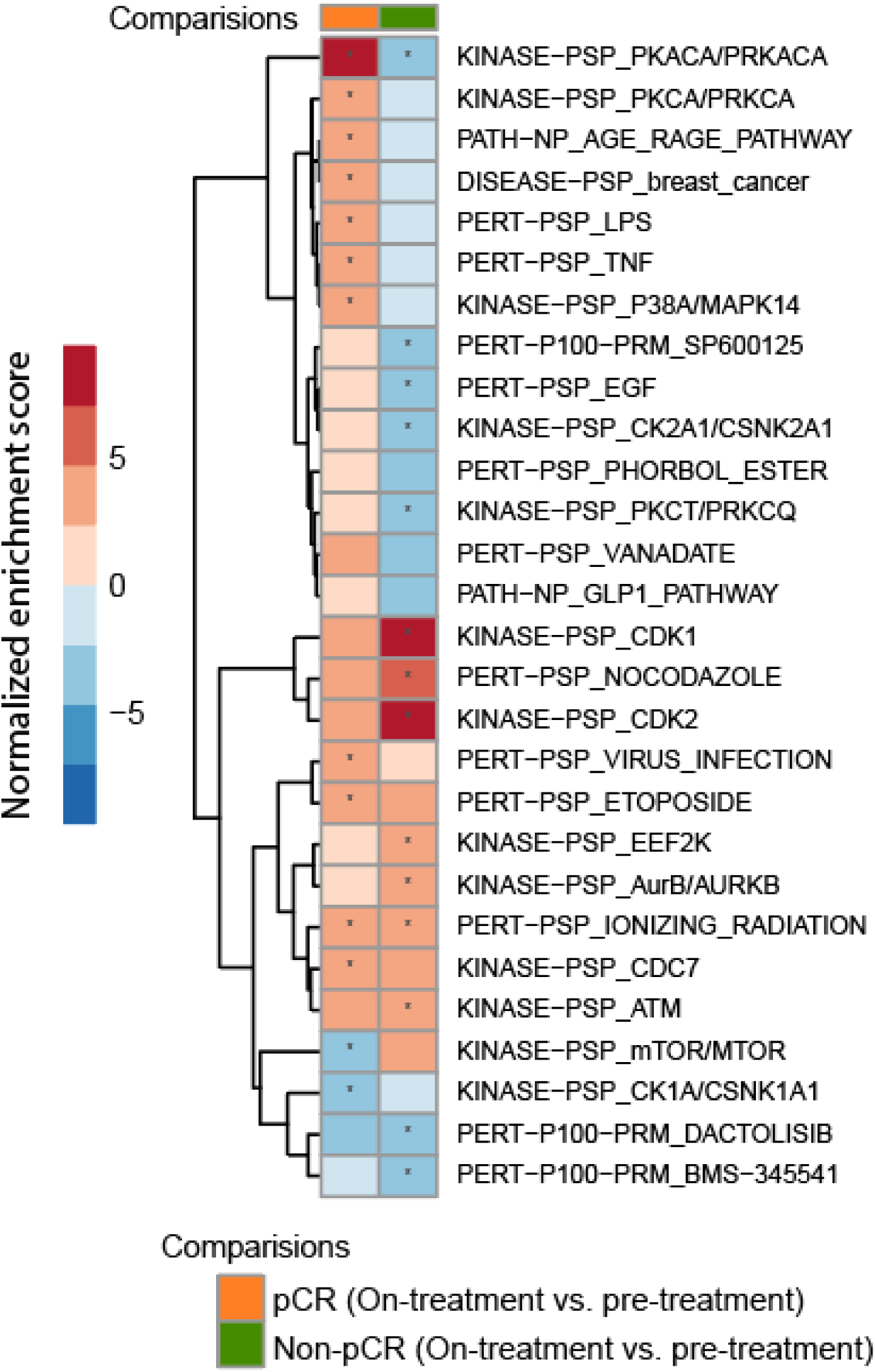
PTM-SEA analysis on pre and on-treatment phosphoproteomics dataset. PTM-SEA was applied to the signed Log10 p-values from *limma* differential expression analysis of on- vs. pre-treatment phosphosite levels from pCR cases (orange) and non-pCR (green) The heatmap shows the Normalized Enrichment Scores (NES) for these kinase signatures, and asterisks indicate significant FDR (<0.05).

**Supplementary Figure 9.**
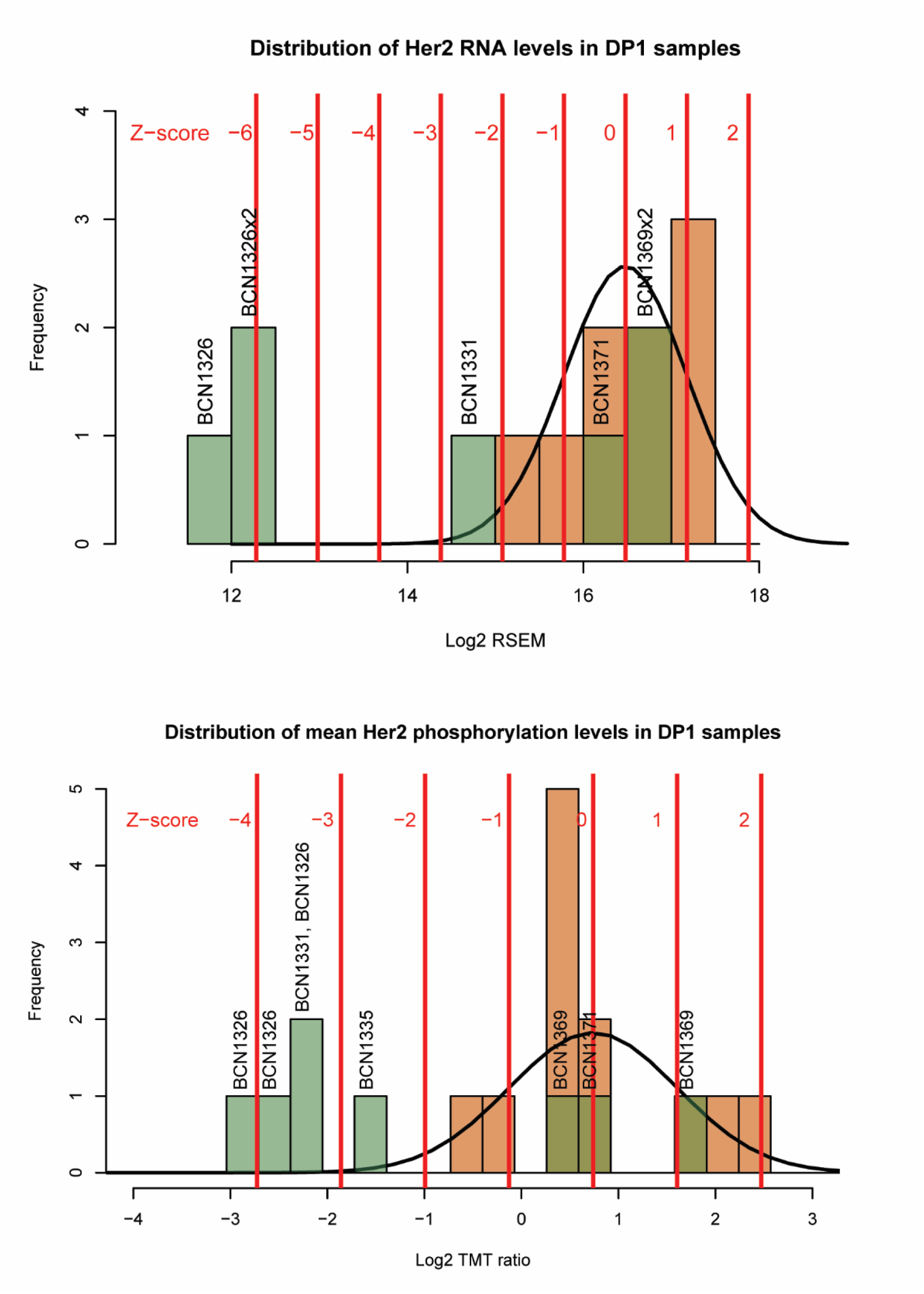
Outlier analysis was performed to identify differentially regulated mRNA, proteins or phosphoproteins in each pre-treatment sample from non-pCR cases relative to the set of pre-treatment samples from all pre-treated pCR cases. Shown is the ERBB2 (HER2) RNA and phosphoprotein distribution across all patients; brown and green bars indicate the frequencies for each protein level bin in non-pCR and pCR, respectively. The line shows the normal distribution of pCR samples from which the Z-score for each non-pCR sample was derived. Z-score thresholds are indicated by red lines.

**Supplementary Figure 10.**
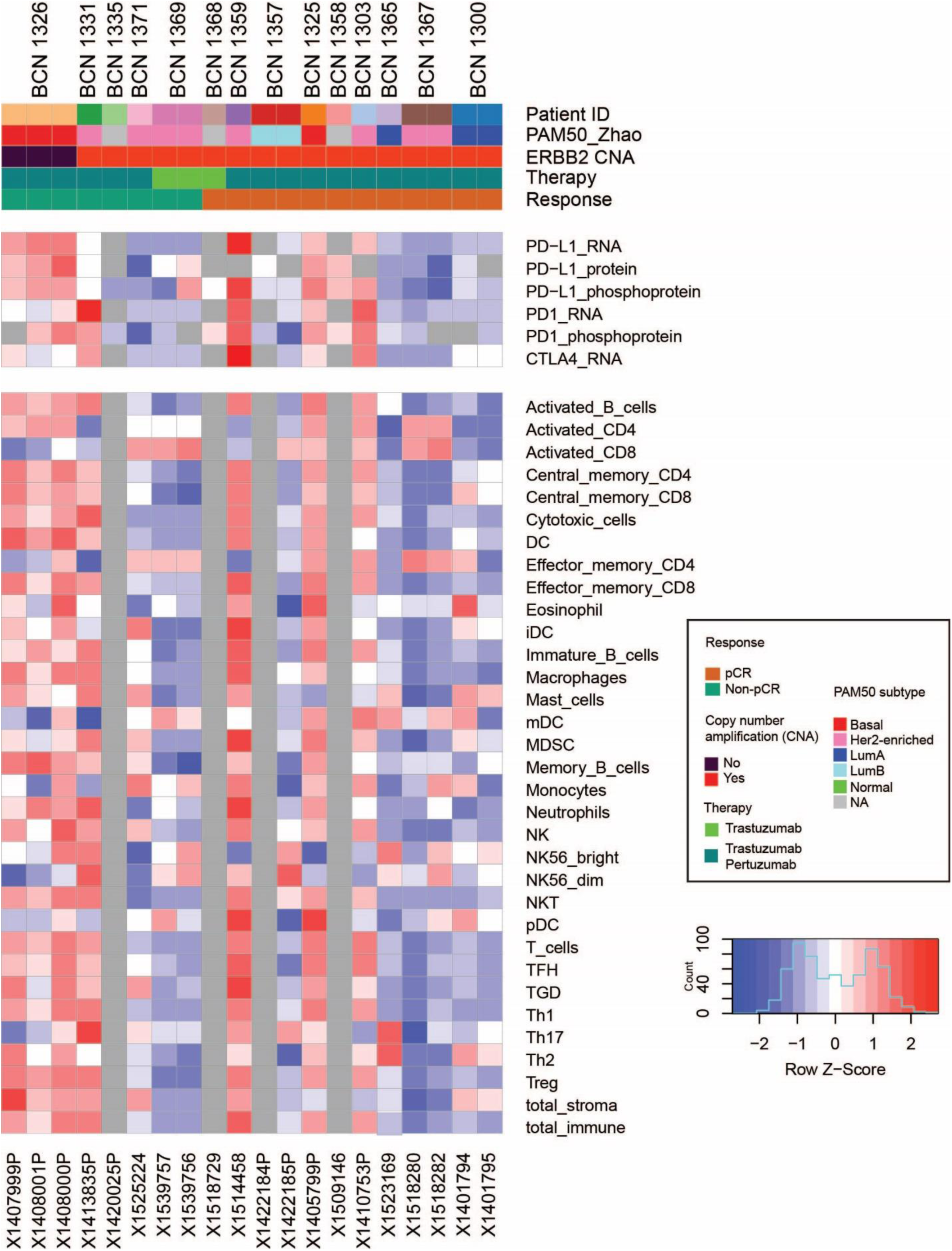
Expression of key immune checkpoint regulators in and immunoprofiling of pre-treated samples. Upper panel shows Z-scores of RNA, protein, and phosphoprotein expression (where available) of key immune checkpoint inhibitors in each baseline sample from pCR (samples on right) and non-pCR (samples on left) patients. Bottom panel shows Z-scores of immune cell profiles inferred from RNA-seq data using Cibersort.

**Supplementary Figure 11.**
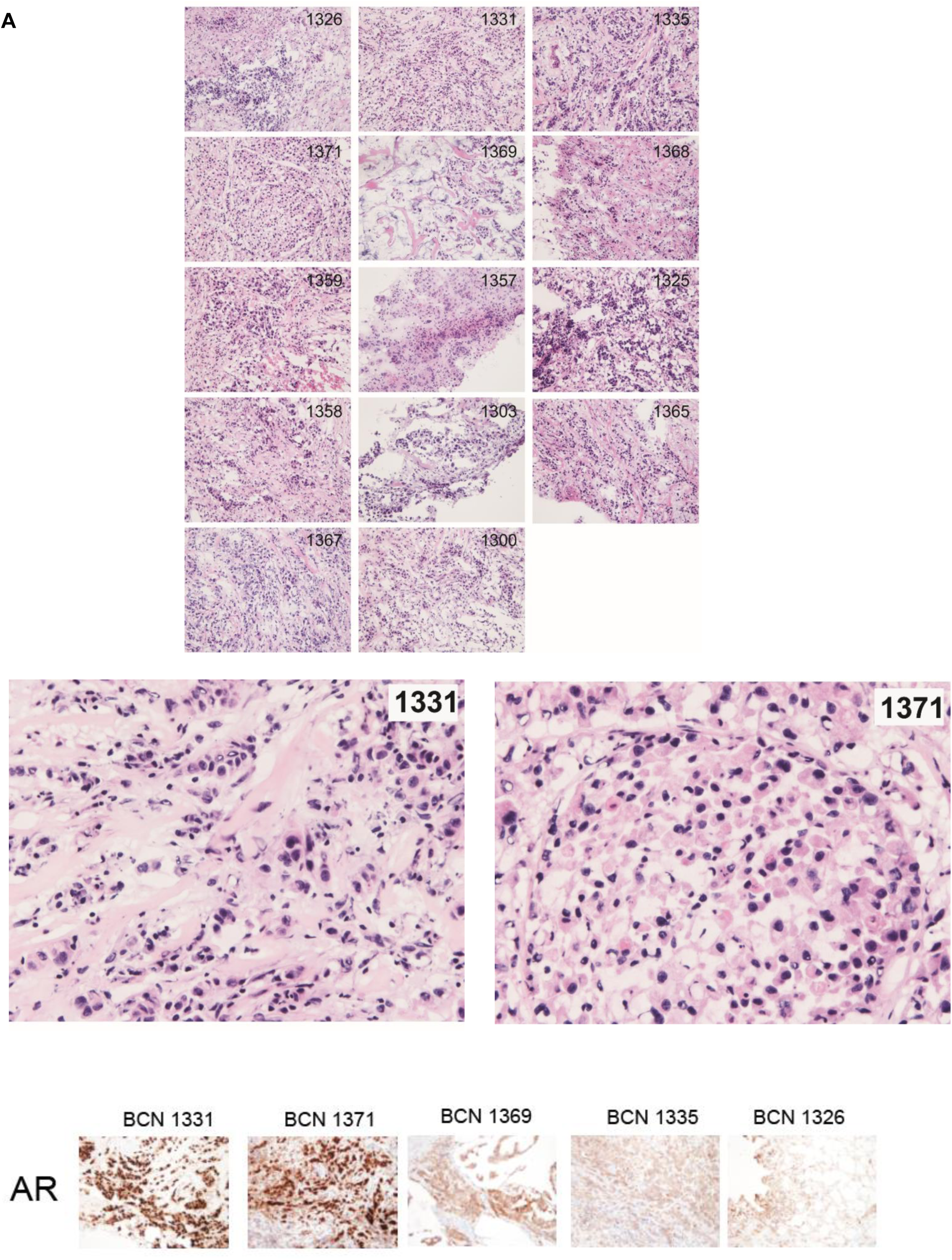
Hematoxylin and eosin (HE) staining of patient samples. HE staining of tissue sections from all 14 patients. The middle panel showed magnified HE stains (400X) of sections patients AR+ patients 1331 and 1371. Patient 1371 shows distinct apocrine features as indicated by plump pink cytoplasm. The lower panel shows immunohistochemical staining profiles of non-pCR cases for AR at x200

**Supplementary Figure 12.**
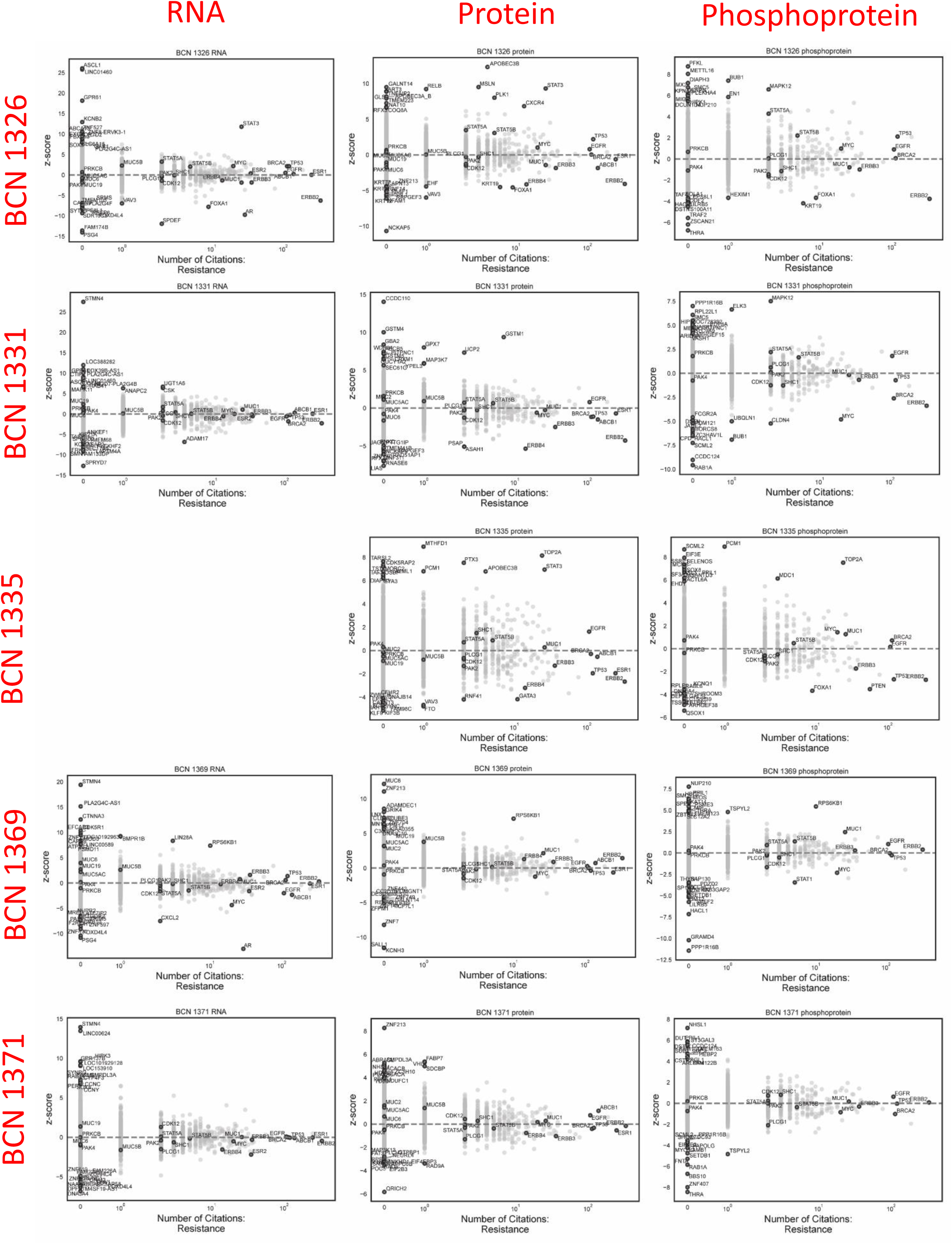
Association of outliers with publications with keywords “breast cancer” and “resistance” or “recur”. Z-score for each gene from outlier analysis is plotted on the y-axis, while the x-axis indicates the number of publications associated with that gene and with breast cancer resistance terms. A separate plot is included for outliers for each non-pCR sample from each omics dataset.

**Supplementary Figure 13.**
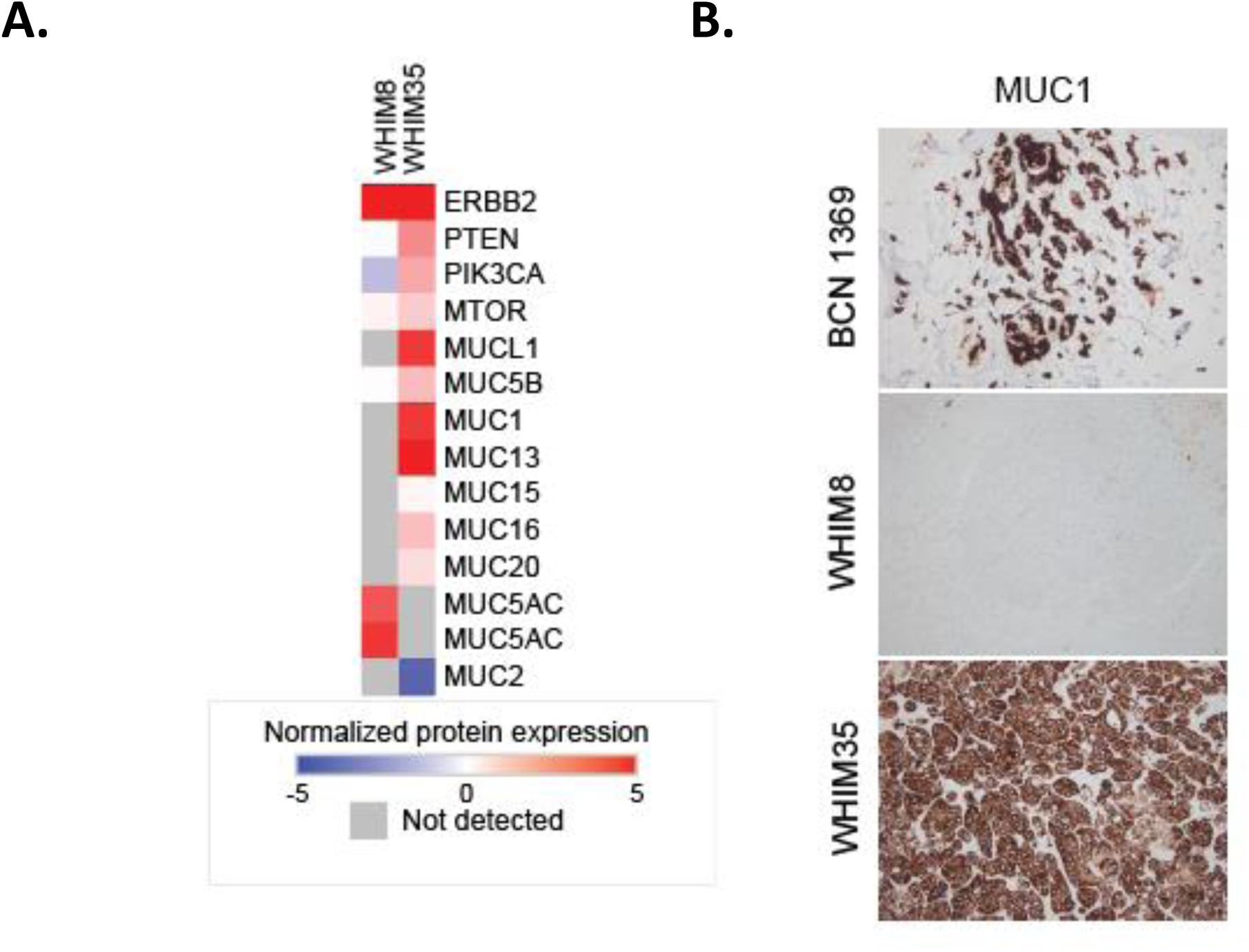
ERBB2 and MUCIN expression in WHIM35, WHIM8 and patient BCN1326 **A**. Heatmap showing ERBB2 pathway and Mucin protein expression in two HER2-enriched PDX (WHIM) models with ERBB2 protein expression. **B**. MUC1 immunohistochemistry (IHC) of WHIM8, WHIM35 and BCN1369.

## Author’s contributions

Conceptualization team: SS, KH, MAG, SAC, MJE; Clinical trial and sample collection: DM, MW, CM, FOA, MR, JH, SJ, GM, MJE; Histopathology team: GM, MJE; Data generation and experimentation: SS, BJ, KH, PS, AG, Lead data analysis: SS, EJ, MEJ, Data analysis: SS, EJ, KK, ABS, MA, CH, NN, YD, BW, BZ, DRM, MAG, SAC, MJE; Intellectual contribution: All authors; Manuscript drafting: SS, EJ, MAG, SAC, MEJ; Manuscript editing: All authors, Funding Acquisition: BZ, DM, MAG, SAC, MEJ.

## Acknowledgments

This work was supported in part by grants from the National Cancer Institute (NCI) Clinical Proteomic Tumor Analysis Consortium grants NIH/NCI U24-CA210986 (to SAC and MAG), NIH/NCI U01 CA214125 (to SAC and MJE), NIH/NCI U24CA210979 (to DRM) and NIH/NCI U24 CA210954 (to BZ). Also, CPRIT established investigator award CPRIT RR140033 to MJE, and NIH/NCI U10 CA180860 (to DM and MJE), NIH/NCI U54CA233223. CPRIT Scholar in Cancer Research was supported by CPRIT Established Investigator Recruitment Award RR140033. Tissue acquisition was also supported by a Breast Cancer Research Foundation (BCRF) grant to MJE. MJE is also a Susan G. Komen Scholar and McNair Medical Foundation Fellow. The authors would like to thank Broad Genomics platform for their assistance with genomic sequencing, Shayan Avanessian and Michael Burgess for technical support and Rena Mao for help with immunohistochemistry. We also thank the Alvin J. Siteman Cancer Center at Washington University School of Medicine and Barnes-Jewish Hospital in St. Louis, MO. and the Institute of Clinical and Translational Sciences (ICTS) at Washington University in St. Louis, for the use of the Tissue Procurement Core, which provided clinical cores. The Siteman Cancer Center is supported in part by an NCI Cancer Center Support Grant #P30 CA091842 and the ICTS is funded by the National Institutes of Health’s NCATS Clinical and Translational Science Award (CTSA) program grant #UL1 TR002345.

## Data availability statement

All genomics and proteomic raw data associated with this study will be available via dbGAP and CPTAC portal (https://proteomics.cancer.gov/data-portal) respectively upon publication.

## Ethical compliance

We have complied with all relevant ethical regulations associated with this study. This study was approved by an institutional review board committee at Broad Institute, Baylor School of Medicine and University of Washington at St. Louis, and the NSABP Foundation. Inc. All patients were consented for proteogenomics analyses.

